# A horizontally transferred fungal deubiquitinase facilitates insecticides resistance in whitefly

**DOI:** 10.64898/2026.06.01.728072

**Authors:** Hantang Lu, Chengjia Zhang, Jinyu Hu, Chao Wang, Rong Zhang, Mingjiao Huang, Qimei Tan, Cheng Yin, Jixing Xia, Yidong Wu, Xuguo Zhou, Ralf Nauen, Youjun Zhang, Chris Bass, Xin Yang

**Affiliations:** State Key Laboratory of Vegetable Biobreeding, Department of Plant Protection, Institute of Vegetables and Flowers, Chinese Academy of Agricultural Sciences, Beijing 100081, PR China; Yuelushan Laboratory, Institute of Plant Protection, Hunan Academy of Agricultural Sciences, Changsha 410125, China; State Key Laboratory of Agricultural and Forestry Biosecurity, Department of Plant Biosecurity, College of Plant Protection, China Agricultural University, Beijing 100193, PR China; College of Plant Protection, Nanjing Agricultural University, Nanjing, PR China; Department of Entomology, School of Integrative Biology, College of Liberal Arts & Sciences, University of Illinois Urbana-Champaign 505 S. Goodwin Ave. Urbana, IL 61801-3795, USA; Crop Science Division, Bayer AG, 40789 Monheim, Germany; Department of Biosciences, College of Life and Environmental Sciences, University of Exeter, Penryn Campus, Penryn, TR10 9FE Cornwall, United Kingdom

**Keywords:** Horizontal gene transfer, Deubiquitylation, Cytochrome P450, *Bemisia tabaci*, metabolism *in vitro*

## Abstract

Horizontal gene transfer (HGT) has enabled insects to acquire novel genetic material that can fuel adaptation to environmental change. However, the role of HGT in the evolution of insecticide resistance remains poorly characterised. Here, we identify *BtUCH19*, a fungal gene that has integrated into the genome of the global pest *Bemisia tabaci* and functions as a deubiquitinating enzyme (DUB). We show that compared to endogenous DUB, BtUCH19 specifically removes K63-linked ubiquitin chains from a cytochrome P450, CYP4C64, thereby stabilizing this key detoxification enzyme *in vivo*. Sustained CYP4C64 abundance enhances metabolic detoxification of two commonly used insecticides thiamethoxam (TMX) and clothianidin (CLO) to their low-toxicity products TMX-Urea and CLO-Urea. Consequently, *BtUCH19* drives insecticide resistance through a novel "HGT-PTM-Metab" axis. Our work reveals the instrumental role of a HGT in orchestrating ubiquitin-proteasome system (UPS)-mediated protein level regulation of a key detoxification pathway. These findings offer new insights into the intricate mechanisms underlying co-evolutionary biology, and provide a new molecular target for Integrated Pest Management (IPM).

## Introduction

Horizontal gene transfer (HGT) enables insects to rapidly acquire novel genetic material that can enhance their environmental adaptability (1). For example, the acquisition of genes by HGT has been shown to mediate adaptation of insect herbivores to host plants by providing a mechanism (enzymatic detoxification) to overcome plant secondary metabolites (2, 3). Recent genomic analysis has suggested that HGT is widespread in insects, and an emerging body of research has revealed roles in various physiological processes including feeding (4, 5), growth, development (6) and reproduction (7), suggesting HGT plays an important role in insect evolution. The globally significant agricultural pest, whitefly *Bemisia tabaci* (Hemiptera: Aleyrodidae), exhibits a high frequency of HGT events (7), and HGT-acquired genes have been implicated in various important physiological processes in this species, including reproduction (8), growth, development (9), and detoxification metabolism (10). Thus, HGT has clearly played an important role in enhancing the adaptability of *B. tabaci,* and in turn its status as an important pest, making it an ideal system to investigate HGT events in insects.

The ubiquitin-proteasome system (UPS) is a vital cellular mechanism that maintains protein homeostasis and thus plays a key role in numerous physiological processes including cell signalling and development, apoptosis and the DNA damage response across eukaryotes (11–14). Emerging evidence suggests that UPS components may also play an important role in the evolution of adaptive traits in arthropods (15, 16). As post-transcriptional modification (PTM) enzymes, deubiquitinating enzymes (DUBs) are important members of the UPS system (17–19), DUBs exert opposing effects to E3 ligases, which specifically recognize ubiquitin molecules linked to substrate proteins and deubiquitinate them, thereby stabilizing these proteins and preventing their degradation by the 26S proteasome (20–22). In insects, DUBs such as USP14 and USP36 regulate various physiological activities, including antiviral responses (23) and antibacterial immunity (24), by stabilizing key metabolic enzymes, representing a novel regulatory layer in insects. Furthermore, certain UPS enzymes are hypothesized to potentially participate in insecticide resistance in insects (15). However, demonstration of the causal role of the UPS system in insect resistance to insecticides remains lacking.

The evolution of insecticide resistance represents a growing challenge to the control of many of the world’s most damaging insect pests. In insects, over five decades of research have demonstrated that insecticide resistance primarily arises from mutations in toxin target sites or changes in metabolic pathways (25–28). In the case of the latter, insect cytochrome P450 enzymes (P450s) have been shown to play a particularly important role in mediating insect resistance to diverse insecticides (29–33). For example, P450s such as CYP6CM1 and CYP4C64 have been shown to mediate resistance to insecticides from multiple classes in *B. tabaci* (34–37). Intriguingly, research has shown that P450s can be subject to ubiquitin / deubiquitin-dependent degradation regulation (38, 39). Considering that DUBs act as antagonists to this process by cleaving ubiquitin chains from P450 proteins, they can regulate P450s post-transcriptionally by preventing their destruction. However, although UPS systems can have a profound impact on various physiological activities of insects, whether DUBs play a role in the regulation of insect P450s and in mediating insecticide resistance, largely remains unknown.

In the current study, we investigated the role of a HGT-acquired deubiquitinase gene in regulating insecticide resistance in *B. tabaci* by its action on the key resistance enzyme CYP4C64. We uncover a unique "HGT-PTM-Meta" detoxification axis that stabilises CYP4C64 via specific K63-deubiquitination to enhance metabolic detoxification, providing new insight into the mechanisms underlying insecticide resistance and the role of HGT in this process.

## Results

### Identification and Characterization of the DUB gene *BtUCH19* in *B. tabaci* Acquired by HGT

A deubiquitinase (DUB) gene, named *UCH19* (Identity: 98.23%, BTA013842.1, GenBank: XM_072302559.1), was identified as significantly upregulated in resistant *Bemisia tabaci* MED (Mediterranean / MED, *SI Appendix*, Fig. S1 and BioProject: PRJNA1401507) compared with susceptible population. The identification of *UCH19* putatively acquired by HGT (horizontal gene transfer) was based on a preliminary survey of *B. tabaci* MED genes involved in specific KEGG (Kyoto Encyclopedia of Genes and Genomes) pathways, a database of HGT genes from Li *et al*. in 2022 (7), and transcriptome sequencing combined with annotation of the *B. tabaci* MED reference genome database (http://www.whiteflygenomics.org/ftp/MED/). *BtUCH19*, is located on the 7th chromosome of *B. tabaci* MED and consists of one exon spanning 2,799 nucleotides, and coding sequence (CDS) spanning 1,176 nucleotides (*SI Appendix*, Fig. S3C). It encodes a protein of 391 amino acids (*SI Appendix*, Fig. S3D) - putatively a ubiquitin carboxyl-terminal hydrolase (*SI Appendix*, Fig. S3A). By cloning and sequencing of *BtUCH19* from *B. tabaci* MED adults (*SI Appendix*, Table S4), we confirmed the accuracy of the genome annotation. Additionally, we selected an endogenous DUBs gene, *BtMYND619*, from the whitefly genome, which exhibits the highest similarity to *BtUCH19* (5.29E^-174^, BTA028619.1, GenBank: XR_011900330.1, *SI Appendix*, Fig. S3A and *Supplementary Material*, Fig. S3A), as a control gene to assist in the study. *BtMYND619* is located on the 8th chromosome, it consists of two exons and one intron (*SI Appendix*, Fig. S3C), with a CDS spanning 600 nucleotides (*SI Appendix*, Fig. S4A), and encodes a protein of 200 amino acids (*SI Appendix*, Fig. S3D). It belongs to the UCH family together with *BtUCH19* (*SI Appendix*, Fig. S3A and B) and also contains a zf-MYND domain (zinc finger MYND, *SI Appendix*, Fig. S3D), which is characteristic of deubiquitinating enzymes.

To confirm that *BtUCH19* is integrated into the *B. tabaci* MED genome, independent genomic analyses were performed. We found that *BtUCH19* is located on scaffold 467 in the MED genome, flanked by the genes *Bta10702* and *Bta10704*. *Bta10702* contains five exons, while *Bta10704* has only one exon, Fig. 1C). Phylogenetic analysis of these two genes showed that they are conserved in insect genomes (*SI Appendix*, Fig. S2 and *Supplementary Material*, Fig. S2). The *B. tabaci* MEAM1 genome also contains *BtUCH19* at this position flanked by the same genes (Fig. 1C). While *BtUCH19* was not found in any other region of the *B. tabaci* genome. The overlapping PCR amplification of those genomic regions confirmed the accuracy of the genome assembly and that *BtUCH19* is indeed integrated into the *B. tabaci* MED genome at this position (Fig. 1D and *SI Appendix*, Table. S6), and the existence of *BtMYND619* was verified in the same way (*SI Appendix*, Fig. S4A). Which strengthens the accuracy that *BtUCH19* is a horizontally transferred gene.

**Fig. 1.**
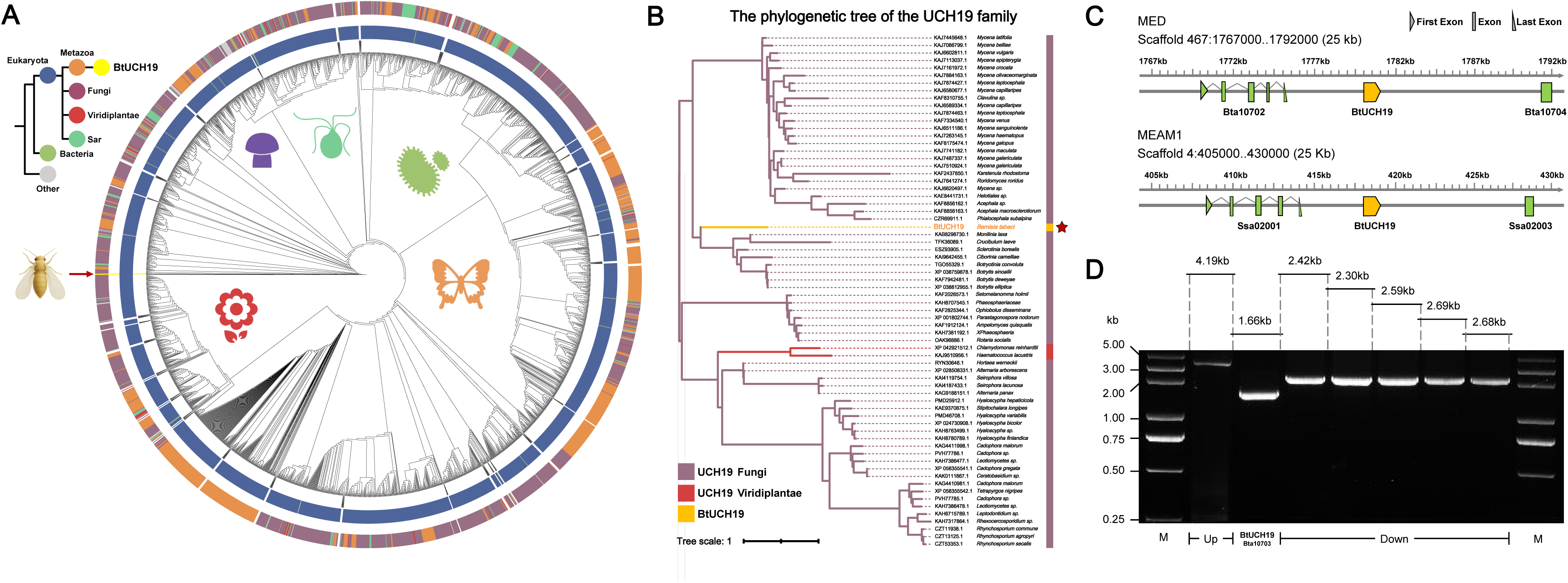
Evidence for the horizontal transfer of *BtUCH19* in *B. tabaci.* (A) Maximum likelihood phylogenetic analyses of *BtUCH19* (CAH0393161.1). *BtUCH19* of *B. tabaci* clusters within a group of fungi ubiquitin carboxyl-terminal hydrolase containing the MYND-type domain. The tree is midpoint rooted, and the scale bar represents 0.5 amino acid substitutions per site. *BtUCH19* is indicated by a red arrow. (**B**) Phylogenetic analysis of *BtUCH19* from various species. The red star indicates *BtUCH19* of the whitefly *B. tabaci* MED. (**C** and **D**) Genome structure of *BtUCH19* and flanking genes, fragments cloned by overlapping PCR from *B. tabaci* MED genome cDNA. Abbreviation: MEAM1: *B. tabaci* Middle East-Asia Minor 1; MED: *B. tabaci* Mediterranean (C). Genome fragment of *BtUCH19* (Bta10703) with its upstream gene (Bta10702) and downstream gene (Bta10704) in MED (D). Note that the phylogenetic tree was constructed using the neighbor-joining algorithm (MEGA11, https://www.megasoftware.net/) from full sequence alignments computed in ClustalX. Bootstrap analyses (1000 replicates) were applied to evaluate the internal support for the tree topology and then visualized via iTOL (https://itol.embl.de/).

To clarify that *BtUCH19* has been acquired by HGT, we compared more than 15,000 protein sequences with similarity to BtUCH19 (XP_072158660.1, *SI Appendix*, Fig. 1A). Investigation of the origin of BtUCH19 using BLASTp and tBLASTn searches against the NCBI non-redundant database revealed that the closest homologs of BtUCH19 are from Fungi (BLASTp E-value < 6.0E-154, bit-score = 548) with no close homologs identified in other arthropods (Fig. 1B). To confirm that *BtUCH19* encodes deubiquitination activity, a deubiquitinase activity assay was used to assess the activity of BtUCH19 *in vivo*, which revealed a significant and time dependent increase in deubiquitinating enzyme (DUB) activity of BtUCH19 compared to the control group (Fig. 2A). Together, these results provide unequivocal evidence that *BtUCH19* encodes a deubiquitinating enzyme that has been horizontally acquired from fungi and integrated into the genome of *B. tabaci* MED.

**Fig. 2.**
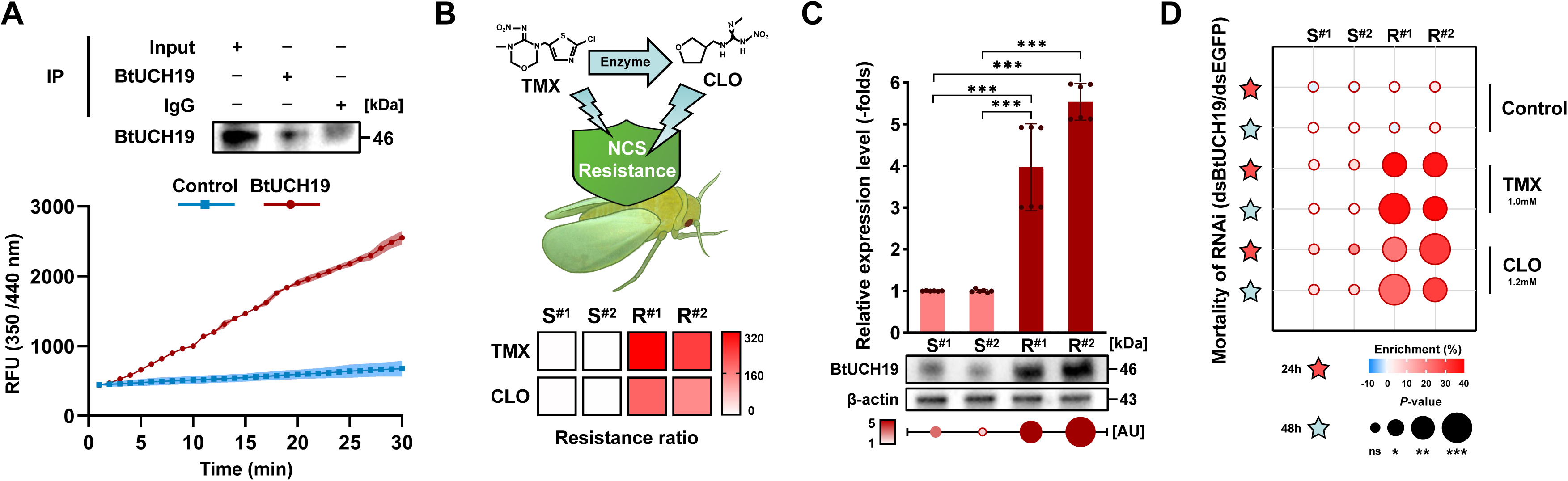
*BtUCH19* is correlated with insecticide resistance in *B. tabaci*. (A) Western blot analyses of BtUCH19 after immunoprecipitation and detection of deubiquitinase activity of BtUCH19 protein in resistant whitefly population R^#2^. An equal volume of 1M PBS was added as the control group for deubiquitinase activity detection (*n* = 3). (B) Determination of resistance ratio (RR) values in comparisons of neonicotinoid-susceptible (S^#1^ and S^#2^) and resistant (R^#1^ and R^#2^) whitefly populations to two neonicotinoids, thiamethoxam (TMX) and clothianidin (CLO). Resistance ratios shown are relative to the reference strain S^#2^. (C) Quantification of *BtUCH19 /* BtUCH19 expression at the mRNA level and protein level among the experimental populations using qPCR and western blot analyses respectively. (D) Sensitivity of the experimental populations to TMX and CLO after RNAi knockdown of *BtUCH19*. Note that all qPCR data are presented as the mean ± SEM of at least three independent experiments. Newly emerged (1-day-old) adults were used in the RNAi experiments and whiteflies fed on ds*EGFP* were used as a negative control. Mortality was assessed 24 h and 48 h after first exposure to insecticide. All bioassay data (mean ± SEM) represent at least three biological replicates with approximately 60 adults in each replication. All western blot analyses represent at least three biological replicates and IgG and *β*-actin were used as loading controls. AU, Relative Intensity of protein. Data were analyzed with ANOVA with Tukey’ s HSD post hoc test: ns (not significant), *P >* 0.05, \**P <* 0.05, \*\**P <* 0.01 and \*\*\**P <* 0.001.

### *BtUCH19* Enhances Resistance to Neonicotinoid Insecticides in *B. tabaci*

Given that *BtUCH19* was upregulated in the transcriptome of a neonicotinoid resistant population of *B. tabaci* (*SI Appendix*, Fig. S1C), we investigated its role in insecticide resistance. First, we examined the expression of *BtUCH19* in other *B. tabaci* populations by collecting one laboratory strain and three field populations of whitefly *B. tabaci* MED, and conducted insecticide bioassays using two neonicotinoid insecticides (thiamethoxam / TMX and clothianidin / CLO). This revealed high levels (144-319-fold) of resistance to both insecticides in resistant populations (R^#1^ and R^#2^) compared with susceptible populations (S^#1^ and S^#2^) in whiteflies (Fig. 2B and *SI Appendix*, Table. S2). Assessment of the messenger RNA (mRNA) levels of *BtUCH19* in these populations revealed an approximately 4-fold higher expression of this gene in resistant strains than in susceptible strains (*P* = 9.36×10^-4^, R^#1^; *P* =1.82×10^-6^, R^#2^, Fig. 2C). However, for *BtMYND619*, only the R^#1^ population exhibits approximately 1.1 times higher expression relative to the S^#1^ strain (*P* = 0.014, *SI Appendix*, Fig. S4B), others show no significant difference in expression. Next, to confirm that overexpression of *BtUCH19* translated to altered expression of the encoded enzyme, we used a rabbit polyclonal antibody raised against a synthetic peptide of BtUCH19 for western blotting to estimate the expression of the encoded enzyme in the susceptible and resistant *B. tabaci* populations. This confirmed that the BtUCH19 protein is expressed at approximately five-times greater levels in resistant populations than in the susceptible populations (Fig. 2C). Then, to demonstrate the causal role of *BtUCH19* in the neonicotinoid resistance of the resistant population, RNA interference (RNAi) was used to knock down the expression of *BtUCH19* (details of synthetic dsRNA can be found in *SI Appendix,* Table. S4). After 6 h of feeding on a diet containing double-stranded RNA (dsRNA) specific for *BtUCH19*, the mRNA levels of *BtUCH19* decreased by 59.8% (R^#2^, *P =* 9.32×10^-8^) and the protein levels of BtUCH19 decreased by approximately two-thirds in R^#2^ (*SI Appendix*, Fig. S5A). Following the reduction of *BtUCH19* / BtUCH19 expression through RNAi treatment, the effect of the mortality of neonicotinoid-treated (bioassay) adults of resistant populations was examined. Knockdown of *BtUCH19* / BtUCH19 significantly increased the mortality of adult whiteflies of the two resistant populations relative to the control (whiteflies fed a diet containing ds*EGFP*: enhanced green fluorescent protein) at 24h and 48h when adult whiteflies were exposed to both a high (Fig. 2D) and low (*SI Appendix*, Fig. S5C) concentration of neonicotinoids (*P* value in *Supplementary Material*, Fig. S5C). However, after interfering with *BtMYND619*, the resistance of the whitefly to pesticides remains almost unchanged (*SI Appendix*, Fig. S4D, *P* value in *Supplementary Material*, Fig. S4D). Collectively, these data provide functional evidence that overexpression of *BtUCH19* enhances resistance to TMX and CLO in *B. tabaci*.

### *BtUCH19* Deubiquitinates Detoxifying Enzymes

To investigate potential substrate proteins of *BtUCH19*, we conducted RNAi by feeding ds*BtUCH19* to adult whiteflies of the resistant population R^#2^, and examined the activity of key superfamilies of detoxification enzymes comprising cytochrome P450 (P450s), glutathione S-transferases (GSTs), and UDP-glycosyltransferases (UGTs). Compared to the control group, knockdown of *BtUCH19* resulted in a 14.45% decrease (*P =* 2.43×10^-2^) in P450 activity, a 14.29% decrease (*P =* 2.63×10^-2^) in GST activity and no significant change (*P =* 4.55×10^-1^) in UGT activity (Fig. 3A). Consequently, we identified P450s and GSTs as potential target proteins for *BtUCH19*’s deubiquitination function. As the P450s and GSTs families of whitefly constitute a large number of genes, we focused on investigation of detoxification enzymes which have been functionally characterized previously (*SI Appendix*, Table. S8). Western blot analysis post-RNAi knockdown of *BtUCH19* identified the P450 enzymes: CYP4C64, CYP306A1, CYP6DW3, CYP4G68, CYP6CM1, and the UGT: UGT354A1 as significantly down-regulated, while the GST: GSTS4 was up-regulated (Fig. 3B, *SI Appendix*, Table. S8), Other detoxifying enzymes, including CYP6EM1, did not show significant changes. Of these, the reduction in the levels of CYP4C64 was particularly pronounced (Fig. 3B), and this P450s has previously been shown to confer resistance to TMX in *B. tabaci* (34). Western blot analysis of CYP4C64 levels after *BtUCH19* knock down in two resistant populations R^#1^ and R^#2^ confirmed these findings (*SI Appendix*, Fig. S4E). For *BtMYND619*, although the expression level of the related mRNA was reduced by 52.05% after treatment with ds*BtMYND619* for 9 hours in R^#1^ (*P* = 0.003, *SI Appendix*, Fig. S4C), the absence of *BtMYND619* has almost no effect on the changes of CYP4C64. All the results above reveal a clear association between *BtUCH19* expression levels and the abundance of CYP4C64, the two variables have a positive correlation. To eliminate interference from other P450 enzymes, we selected a P450 detoxification enzyme, CYP6EM1 (33), which is highly expressed in the resistant population but not affected by BtUCH19 (*SI Appendix*, Fig. S5H), to be used as a control P450s in subsequent experiments.

**Fig. 3.**
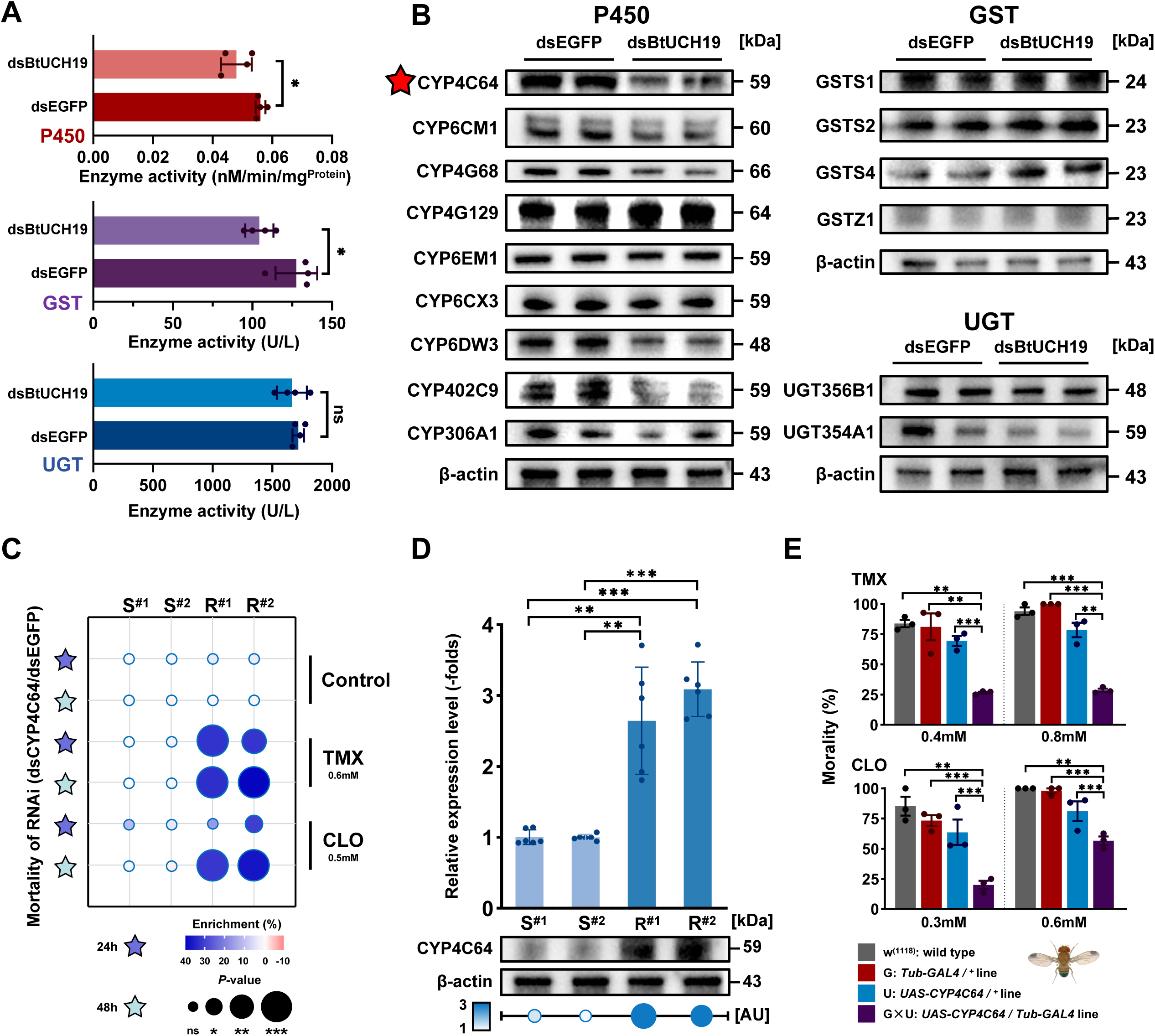
*BtUCH19* is involved in detoxification of neonicotinoids by regulating the cytochrome CYP4C64. (A) The effect of RNAi knockdown of *BtUCH19* on P450, GST and UGT total activity in resistant population R^#2^. (B) The abundance of key detoxification enzymes in *B. tabaci* following RNAi. Western blot to detect the abundance of related detoxification enzyme proteins after ds*BtUCH19* RNAi in R^#2^. (C) Sensitivity of the experimental populations to thiamethoxam (TMX) and clothianidin (CLO) after RNAi knockdown of *CYP4C64*. (D) Quantification of the expression of the mRNA and protein levels of *CYP4C64 /* CYP4C64 among the experimental populations using qPCR and western blot analyses, respectively. (E) Sensitivity of transgenic *Drosophila* expressing *CYP4C64* to TMX and CLO compared to the wild-type line and the parental line that does not actively express the transgene; *w^(1118)^*, wild type; G, *Tub-GAL4/^+^* line; U, *UAS-CYP4C64/^+^* line; G×U, *UAS-CYP4C64/Tub-GAL4* line). Note that all qPCR data are presented as the mean ± SEM of at least three independent experiments. Newly emerged (1-day-old) adults were used in the RNAi experiments and whiteflies fed on ds*EGFP* were used as a negative control. Mortality was assessed 24 h and 48 h after first exposure to insecticide. All bioassay data (mean ± SEM) represent at least three biological replicates with approximately 60 adults in each replication. All western blot analyses represent at least three biological replicates and *β*-actin was used as a loading control. AU, Relative Intensity of protein. Data were analyzed with ANOVA with Tukey’ s HSD post hoc test: ns = not significant, *P >* 0.05, \**P <* 0.05, \*\**P <* 0.01 and \*\*\**P <* 0.001.

### *CYP4C64* Confers Resistance to Neonicotinoids *In Vivo*

To verify the role of CYP4C64 in neonicotinoid resistance in our four experimental populations of *B. tabaci*, we first quantified *CYP4C64 /* CYP4C64 expression in the experimental populations using real-time quantitative polymerase chain reaction (qPCR) and western blot analysis. The results showed that *CYP4C64 /* CYP4C64 was significantly upregulated at both the mRNA and protein levels in resistance compared with susceptible (Fig. 3D). Specifically, for *CYP6C64* / CYP6C64, the mRNA expression in resistant populations is three times that of susceptible populations (*P* = 2.78×10^-3^, R^#1^; *P* = 2.37×10^-5^, R^#2^ respectively), and the protein levels in resistant populations is 3 times that of susceptible populations. After treating adult resistant whiteflies of the R^#2^ strain with ds*CYP4C64* for 36 hours, the expression of *CYP4C64* significantly decreased 65.77% (*P =* 9.93×10^-7^) compared to the control (fed ds*EGFP*). The corresponding protein levels decreased by approximately half (*SI Appendix*, Fig. S5B). Knockdown of *CYP4C64* / CYP6C64 significantly increased the mortality of adult whiteflies of the two resistant populations, relative to the control (fed ds*EGFP*) at 24h and 48h when exposed to both a low (Fig. 3C) and a high (Fig. S5F) concentration of two neonicotinoids (*P* value in *Supplementary Material*, Fig. S5F).

To provide additional evidence that *CYP4C64* confers tolerance to neonicotinoids *in vivo*, we used wild type and transgenic *Drosophila melanogaster* lines to validate the function of this P450 gene. qPCR and western blot analyses demonstrated that transgenic *Drosophila* expressing *CYP4C64* (G×U: *UAS-CYP4C64/Tub-GAL4* line) exhibited higher levels of *CYP4C64 /* CYP4C64 expression compared with the wild-type *w^(1118)^*, parental line (G: *Tub-GAL4/^+^* line) and the parental line that does not actively express the transgene (U: *UAS-CYP4C64/^+^* line). The G×U line exhibited a higher *CYP4C64* mRNA level, approximately 23.9-fold higher than that of *w^(1118)^* line (*P* = 1.81×10^-11^), 4.7-fold higher (*P* = 1.71×10^-10^) than that of the G line and 1.9-fold higher (*P* = 3.32×10^-9^) than that of the U line (*SI Appendix*, Fig. S5G), the protein expression level in the G×U line was approximately 20 times higher than that in the *w^(1118)^*, G and U lines (*SI Appendix*, Fig. S5J). Insecticide bioassays showed that the G×U line exhibits significant resistance to two neonicotinoids compared with the *w^(1118)^*, G line and U line (Fig. 3E, *P* value in *Supplementary Material*). Consistent with previous work (34), these findings provide clear evidence that *CYP4C64* confers resistance to two neonicotinoids *in vivo*.

### CYP4C64 Metabolism of Neonicotinoids

To demonstrate that CYP4C64 can detoxify neonicotinoids directly, the baculovirus-mediated expression system was employed to heterologously express CYP4C64 *in vitro*. Expression of functional recombinant P450s was confirmed by reduced CO-difference spectroscopy, revealing a characteristic absorbance peak at 450 nm indicative of catalytically active P450s. And western blotting was also used to assess expression levels in H5 (high five) insect cells (Fig. 4A).

**Fig. 4.**
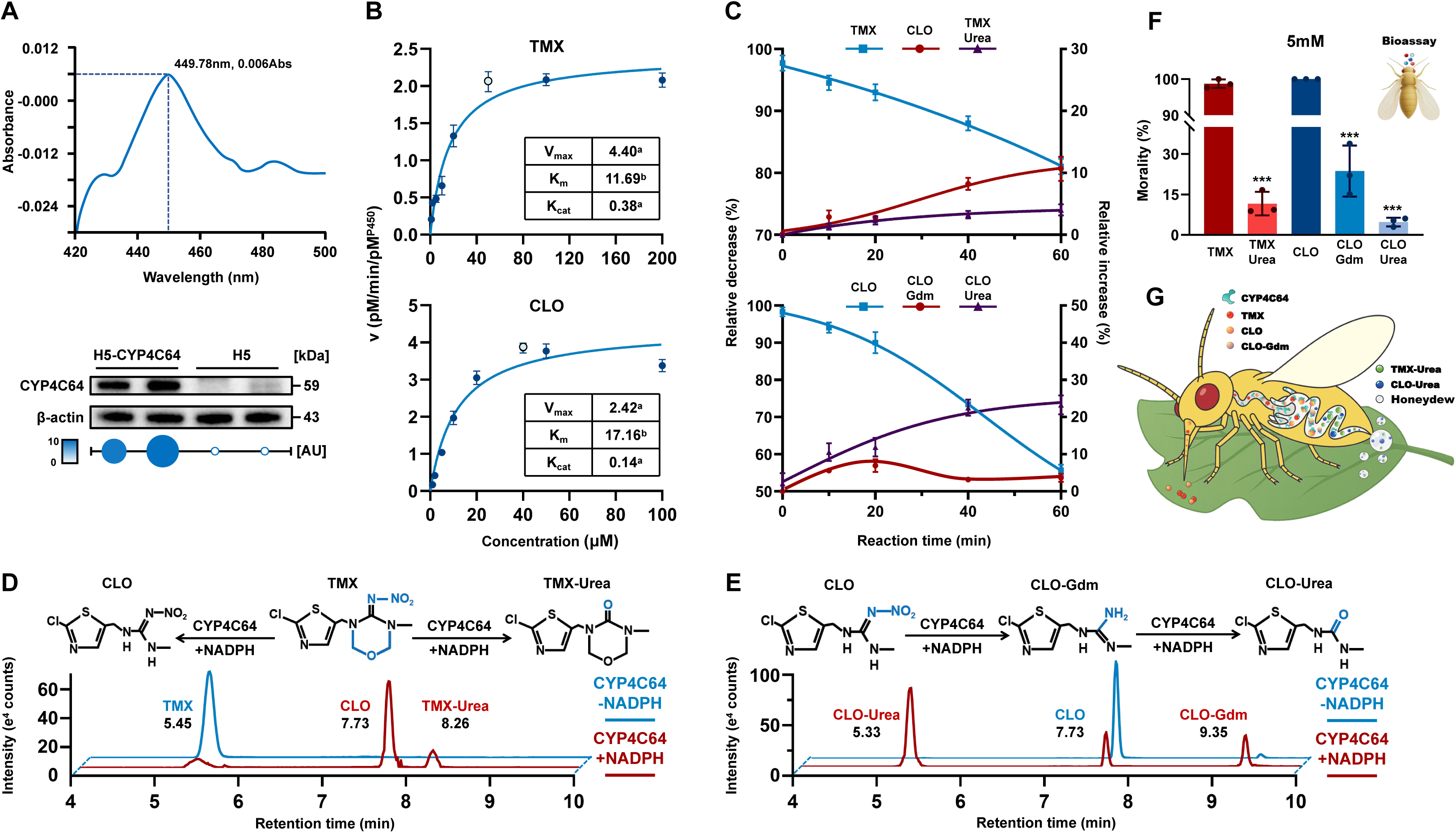
CYP4C64-mediated metabolism of neonicotinoid insecticides. (A) Reduced CO-difference spectra of recombinant CYP4C64 and western blot analysis of the levels of CYP4C64 in empty viral plasmids, H5 cells and P450-expressing H5 cells. (B) Michaelis-Menten kinetics of thiamethoxam (TMX) and clothianidin (CLO) metabolism by CYP4C64 (the hollow points indicate the substrate optimum concentrations for detecting the metabolic capability of CYP4C64. V_max_, Maximum rate of metabolism; K_m_, Michaelis constant; K_cat_, Catalytic constant: V_max_ / K_m_; a, pmol / min / pmol^P450^; b, μM. Curves were generated by fitting the Michaelis-Menten equation to determine CYP4C64 enzyme kinetic constants. (C) Time course of TMX and CLO depletion and metabolites (CLO, thiamethoxam-Urea / TMX-Urea, clothianidin-guanidine / CLO-Gdm and clothianidin-Urea / CLO-Urea), Points are fitted to Sigmoidal, 4PL. Reactions were performed at 30°C with 50 pM of TMX and 40 pM of TMX. (D and E) UPLC–MS analyses of TMX and CLO metabolites. Multi-Reaction-Monitoring (MRM) chromatograms of CYP4C64 catalysed TMX (D) and CLO (E) metabolism: Red, P450 microsomes incubation of 2 nmol neonicotinoid for 60 min with NADPH; Blue, P450 microsomes incubation without NADPH. The molecular formula above indicates the production of metabolic products and changes in chemical bonds during the process. (F) The sensitivity of resistant population R^#2^ to 5 mM TMX, CLO, TMX-Urea, CLO-Gdm and CLO-Urea. (G) Figure illustrating the process by which TMX and CLO undergo detoxification after being metabolized by CYP4C64. Note that CO-difference spectral tests and all UPLC–MS analyses data are presented in at least three independent experiments. All UPLC-MS analyses data are presented from at least three independent experiments. Mass concentrations were calculated using relative standard curves, with microsomes incubated without NADPH serving as the control. All bioassay data (mean ± SEM) represent at least three biological replicates with approximately 60 newly emerged (1-day-old) adults in each replication. Western blot experiments comprised at least three biological replicates and *β*-actin was used as a loading control. AU, Relative Intensity of protein. Data were analyzed with ANOVA with Tukey’ s HSD post hoc test: \*\*\**P <* 0.001.

To clarify the metabolic capacity of mature CYP4C64 protein towards two neonicotinoids, the abundance of neonicotinoids was evaluated after incubation with CYP4C64 through chromatographic peak intensity quantification using ultra-performance liquid chromatography-mass spectrometry (UPLC-MS) analysis. Comparing with NADPH-depleted samples as the control and using the corresponding substance’s standard curve, under conditions of sufficient CYP4C64, the concentration of substrate insecticides in NADPH-supplemented reaction systems revealed a 18.27% reduction (9.23 pmol, *P* = 1.37×10^-3^) in TMX and a 53.29% reduction (17.73 pmol, *P* = 2.46×10^-6^) in CLO (*SI Appendix*, Fig. S6A) following 1h incubation of recombinant protein and insecticide. The CYP4C64 metabolism of neonicotinoids followed Michaelis-Menten kinetics; for TMX, maximum metabolic rate (V_max_) = 2.42 pmol / min / pmol^P450^, Michaelis constant (K_m_) = 17.16 μM, catalytic efficiency (K_cat_, V_max_ / K_m_) = 0.14 pmol / min / pmol^P450^; for CLO, V_max_ = 4.40 pmol / min / pmol^P450^, K_m_ = 11.69 μM, K_cat_ = 0.38 pmol / min / pmol^P450^ (The number of testing objects / N, degrees of freedom / Df and coefficient of determination / R^2^ of CYP4C64 toward TMX and CLO are listed in Fig. 4B and *SI Appendix*, Table. S9). Compared to TMX, CYP4C64 exhibits a higher V_max_ and lower K_m_ toward CLO, resulting in approximately 3-fold greater K_cat_ for CLO. Based on protein concentration normalization via Bradford assay and cytochrome P450 heme quantification, the enzyme-specific activity was calculated as 145.2 pM of TMX or 264.0 pM of CLO metabolized per pmol of CYP4C64 protein in 60 minutes.

To investigate the metabolic pathways of TMX and CLO following metabolism by CYP4C64, we analyzed their specific metabolic products using UPLC-MS. We identified the characteristic mass spectrometry peaks of TMX (eluting at 6.43 min) and CLO (eluting at 7.78 min) as substrates participating in the reaction (*SI Appendix*, Fig. S6B and Table. S11) and several characteristic mass spectrometry peaks of their potential metabolic products (*SI Appendix*, Fig. S6C and Table. S11). By comparing the retention times of potential metabolic products and reference standards in UPLC-MS, under the catalysis of CYP4C64, TMX is converted into two primary metabolites: CLO (retention time at 7.73 min) and thiamethoxam-urea (TMX-Urea, retention time at 8.26 min) respectively (Fig. 4D); CLO is transformed into two main metabolites, which were clothianidin-guanidine (CLO-Gdm, retention time at 9.35 min) and clothianidin-urea (CLO-Urea, retention time at 6.79 min) respectively (Fig. 4E). TMX and CLO depletion and metabolite peak formation were time-dependent (Fig. 4C). Within 60 min in the CYP4C64+NADPH environment, TMX decreased by 18.97% (*P* = 1.21×10^-3^), the product CLO in the system increased by 10.67% (*P* = 1.01×10^-2^), and the product TMX-Urea increased by 4.03% (*P* = 1.64×10^-2^). CLO, as the substrate of the enzymatic reaction, decreased to 44.33% (*P* = 1.96×10^-4^), the product CLO-Gdm first increased and then decreased, with a peak of 6.92% (*P* = 2.01×10^-2^), and the product CLO-Urea increased by 23.28% (*P* = 2.32×10^-3^). These results illustrate the dynamic changes in substrate and products in the metabolism of TMX and CLO mediated by CYP4C64.

### Toxicological Analysis of Neonicotinoid Metabolites

To confirm that these compounds are the final metabolites produced by CYP4C64 metabolism, we conducted substrate / metabolite bioassays on resistant *B. tabaci* population R^#2^ using two different concentrations of the compounds, and collected the honeydew of whiteflies for metabolic analyses. Compared to TMX and CLO, the toxicity of their metabolites to *B. tabaci* was significantly reduced, with the exception of CLO produced upon TMX metabolism as previously reported (40-42), CLO exhibiting greater toxicity than TMX at equal concentration (Fig. 2B and *SI Appendix*, Table. S1). For other TMX metabolites, at a concentration of 5 mM, the mortality rate of *B. tabaci* fed TMX-Urea was 11.76% lower (*P* = 4.92×10^-6^) than those fed TMX; The mortality rate of *B. tabaci* fed CLO-Gdm was 23.69% lower (*P* = 1.52×10^-4^) than those fed CLO; the mortality rate of *B. tabaci* fed CLO-Urea was 4.78% lower (*P* = 5.75×10^-8^) than those fed CLO (Fig. 4F). These findings were consistent using an administration concentration of 3 mM (*SI Appendix*, Fig. S7C). Finally, we fed the whiteflies with a diet solution containing two insecticides (50 mg/L, respectively) after interfering with *CYP4C64*, and detected the components in honeydew after 48 h in R^#2^ (*SI Appendix*, Fig. S7A). The UPLC-MS analysis of honeydew showed that, there are two metabolites, TMX-Urea and CLO-Urea (*SI Appendix*, Fig. S7A). Compared with the control group (ds*EGFP*), after interfering with *CYP4C64* / *BtUCH19* / *TRIM37* / *BtMYND619*, the two main metabolites in the honeydew were significantly reduced following the loss of CYP4C64 / BtUCH19 (for CYP4C64: TMX-Urea, 50.52%, *P* = 0.0048, CLO-Urea, 38.82%, *P* = 0.022; for BtUCH19: TMX-Urea, 32.59%, *P* = 0.016; CLO-Urea, 44.71%, *P* = 0.011; *SI Appendix*, Fig. S7D), while the deletion of TRIM37 resulted in increased levels of TMX-Urea and CLO-Urea in the honeydew (TMX-Urea, 41.49%, *P* = 0.022; CLO-Urea, 29.18%, *P* = 0.042; *SI Appendix*, Fig. S7D). This suggests that the absence of CYP4C64 and BtUCH19 weakens whiteflies’ metabolic capacity toward two neonicotinoids, whereas the loss of TRIM37 enhances the tendency of TMX and CLO to convert into urea products. Additionally, for the control gene *BtMYND619* of *BtUCH1*9, its deletion had almost no effect on the metabolism of TMX and CLO (*SI Appendix*, Fig. S7D). As presented above, both TMX-Urea and CLO-Urea exhibited lower toxicity than their respective parent compounds and demonstrate increased hydrophilicity, which enhances excretion (Fig. 4G). Thus, CYP4C64 completes the detoxification process of neonicotinoids by metabolizing hydrophobic substrates into more hydrophilic products that contain a urea group, making them less toxic to *B. tabaci*.

### *TRIM37* and *BtUCH19* act in concert to regulate CYP4C64

Given that deubiquitination and ubiquitination are dynamic reversible processes, we next investigated the role of the ubiquination gene *TRIM37* in the regulation of CYP4C64 in *B. tabaci*. We identified TRIM37 as an essential ubiquitin ligase for E3 activity and revealed that it is down-regulated in *B. tabaci* resistant populations based on our transcriptome sequencing results (Fig. S1 and *SI Appendix*, Fig. S1). The result demonstrated that the mRNA expression of *TRIM37* in *B. tabaci* resistant populations was lower (16.65%) than in susceptible populations (*P* = 6.49×10^-9^, S^#1^; *P* = 4.98×10^-9^, S^#2^ respectively), and the protein expression of TRIM37 in resistant populations was half of that in susceptible populations (Fig. 5A). We then used RNAi to knock down *TRIM37* and measured expression levels of *TRIM37/*TRIM37 and CYP4C64. After treating the susceptible population S^#2^ with ds*TRIM37* for 24 hours, the expression of *TRIM37 /* TRIM37 significantly decreased by 55.81% at the mRNA (S^#2^, *P =* 2.05×10^-8^, *SI Appendix*, Fig. S5D) and by approximately half at the protein level (Fig. 5B), compared to the control (fed ds*EGFP*). In contrast, the protein expression of CYP4C64 increased approximately three-fold (S^#1^ and S^#2^, Fig. 5B). In addition, knockdown of *TRIM37* significantly increased the mortality of susceptible whiteflies, relative to the control (fed ds*EGFP*), at 24h and 48h when adult whiteflies were exposed to both a low (Fig. 5C) and high (*SI Appendix*, Fig. S5I) concentration of a range of neonicotinoids (*P* value in *Supplementary Material*, Fig. S5I). These findings suggest that *TRIM37* exerts a negative regulatory effect on the abundance of CYP4C64. Moreover, the low expression of *TRTIM37* is unfavorable for the development of resistance in *B. tabaci*.

**Fig. 5.**
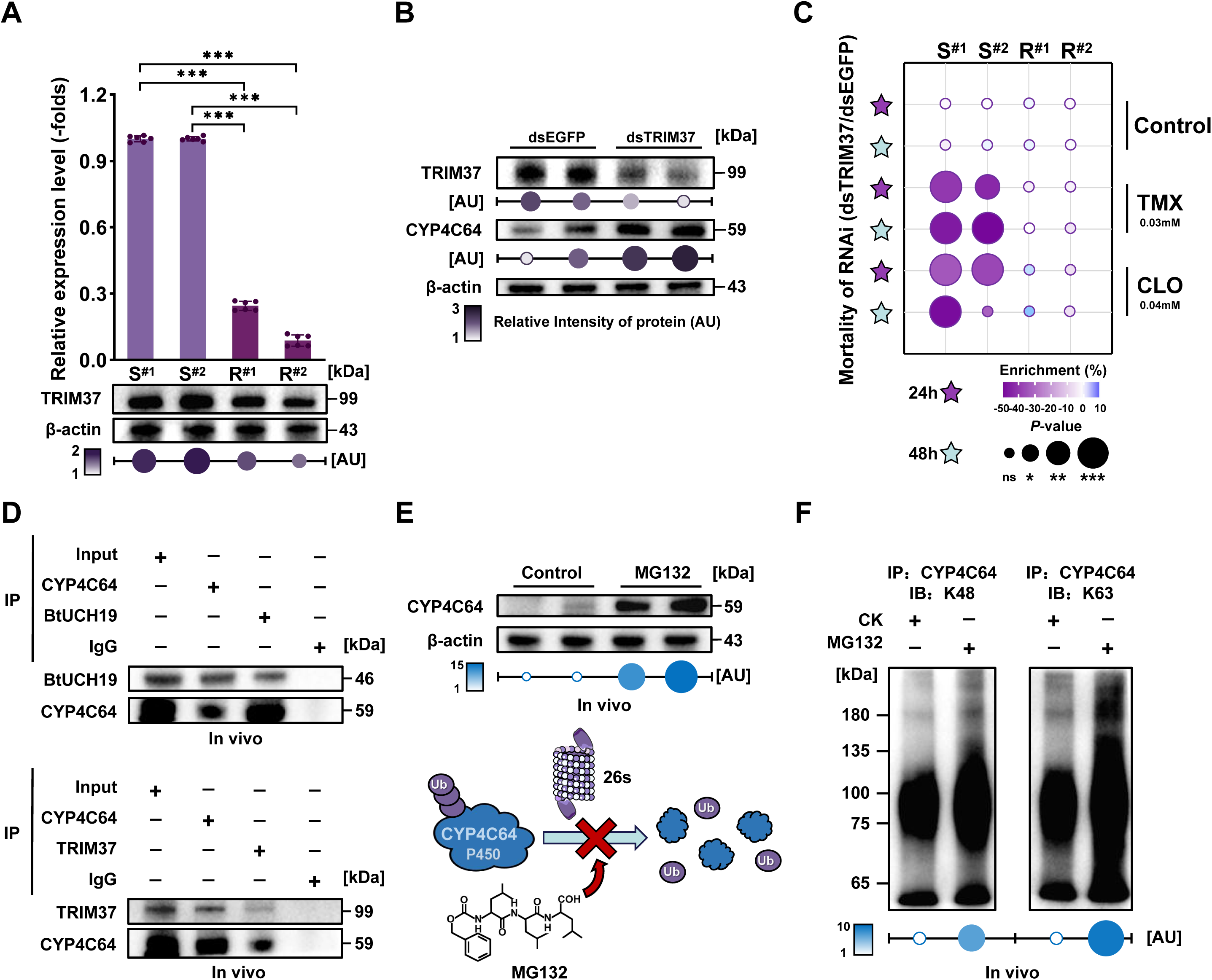
*BtUCH19* and *TRIM37* jointly regulate the ubiquitin-proteasome degradation pathway of CYP4C64. (A) Quantification of the expression of the mRNA level and protein levels of *TRIM37 /* TRIM37 among the experimental populations as assessed by qPCR and western blot. (B) Western blot analyses of the expression of TRIM37 and CYP4C64 after RNAi knockdown of *TRIM37* in two susceptible populations (S^#1^ and S^#2^). (C) Sensitivity of the experimental populations to thiamethoxam (TMX) and clothianidin (CLO) after RNAi knockdown of *TRIM37*. (D) Immunoprecipitation (IP) analyses of interaction of BtUCH19 / TRIM37 with CYP4C64 in the resistant population R^#2^. The extracted protein from whiteflies was incubated with the antibody of BtUCH19 / TRIM37 and CYP4C64, followed by incubation with protein A / G-magnetic beads. Then the BtUCH19 / TRIM37 and CYP4C64 proteins were detected by their corresponding antibodies in western blots. (E) CYP4C64 protein level *in vivo* after being fed with the 26S proteasome inhibitor MG132 (50 mM) in the S^#2^, with DMSO used as a control (CK). The CYP4C64 protein was detected by anti-CYP4C64 in western blots. (F) The degradation of CYP4C64 via the 26S proteasome pathway in the R^#2^. The total protein was extracted from whiteflies fed with MG132 (50 mM) for 24 h, with DMSO used as a control. Then the protein was immunoprecipitated with anti-CYP4C64 and detected by anti-K48 / anti-K63 in western blots. Note that all qPCR data are presented as the mean ± SEM of at least three independent experiments. Newly emerged (1-day-old) adults were used in the RNAi experiments and whiteflies fed on ds*EGFP* were used as a negative control. Mortality was assessed 24 h and 48 h after the first exposure to insecticides. All bioassay data (mean ± SEM) represent at least three biological replicates with approximately 60 adults in each replication. All western blot analyses represent at least three biological replicates and Input, IgG (immunoglobulin G) and β-actin were used as loading controls. AU, Relative Intensity of protein. Data were analyzed with ANOVA with Tukey’s HSD post hoc test: ns (not significant), P > 0.05, *P < 0.05, **P < 0.01 and ***P < 0.001.

After identifying that *BtUCH19* and *TRIM37* have opposing regulatory effects on CYP4C64, we used co-immunoprecipitation (co-IP) to further investigate whether CYP4C64 is a substrate P450 of BtUCH19 and TRIM37. In the co-IP assay, antibodies against BtUCH19 and TRIM37 were used to detect total protein from whitefly R^#2^ incubated with CYP4C64 antibody with signals detected for both BtUCH19 and TRIM37 (Fig. 5D), indicating that BtUCH19 and TRIM37 can directly interact with CYP4C64 *in vivo*. When treated in the same manner, CYP6EM1 was found not to directly bind to BtUCH19 or TRIM37 (*SI Appendix*, Fig. S9A), which excludes false-positive results for CYP4C64.

To confirm that BtUCH19 deubiquitinates CYP4C64, and if TRIM37 mediates CYP4C64 ubiquitination triggering its degradation, CYP4C64 protein was detected via western blotting following treatment with MG132, a potent inhibitor of degradation of ubiquitin-conjugated proteins by the 26S proteasome (43) treated with DMSO served as the control group. The presence of MG132 resulted in an approximately 15-fold increase in the abundance of CYP4C64 compared with the control group (Fig. 5E). The results indicated that when the ubiquitin-proteasome pathway was inhibited, the degradation of CYP4C64 was suppressed. Next, to clarify the specific pathway by which CYP4C64 participates in the UPS (ubiquitin-proteasome system), western blots were used to detect CYP4C64 using K48 and K63 ubiquitin antibodies. The abundance of the K48 pathway increased approximately 5-fold, while the abundance of the K63 pathway increased more than 10-fold (Fig. 5F), which suggested that the K63 pathway plays a more prominent role in degrading CYP4C64 *in vivo.* In summary, these data reveal that both BtUCH19 and TRIM37 can bind to CYP4C64 directly and guide it to participate in the UPS, and that CYP4C64 is degraded primarily via the K63 pathway.

### *BtUCH19* Deconjugates the K63-Linked Ubiquitin Chain from *CYP4C64* at Lys236

To determine the structural domains involved in the interaction between BtUCH19 / TRIM37 and CYP4C64, we predicted multiple ubiquitination sites of CYP4C64 with RUBI Version 1.0 (http://old.protein.bio.unipd.it/rubi/). Then the wild-type vectors of BtUCH19 / TRIM37 and CYP4C64, and the mutation vectors CYP4C64^162^, CYP4C64^164^, CYP4C64^198^, CYP4C64^236^, and CYP4C64^281^, where the lysine / K residue at these positions was mutated to arginine / R (*SI Appendix*, Table. S7), were generated *in vitro* and yeast two-hybrid assays (Y2H) run to explore interactions between these constructs (*SI Appendix*, Table. S7). Using BtMYND619 as the control group and excluding false-positive results (*SI Appendix*, Fig. S9B), TRIM37 and BtUCH19 were found to directly interact with CYP4C64, consistent with our earlier results, the mutation vectors CYP4C64^162^ and CYP4C64^236^ did not strongly interact with either TRIM37 or BtUCH19 (Fig. 6A and *SI Appendix*, Table. S5B). Notably, while the mutation construct CYP4C64^162^ exhibits weak interaction with TRIM37, the CYP4C64^236^ construct showed almost no interaction with either BtUCH19 and TRIM37, especially in quadruple dropout synthetic dextrose medium (SD, -Ade / Adenine, -His / Histidine, -Leu / Leucine, - Trp / Tryptophan). Thus, Y2H experiments provide initial evidence that BtUCH19 and TRIM37 interact with CYP4C64 *in vitro*, and that ubiquitination sites at K residues 162 and 236 of CYP4C64 may be important binding sites in this process.

**Fig. 6.**
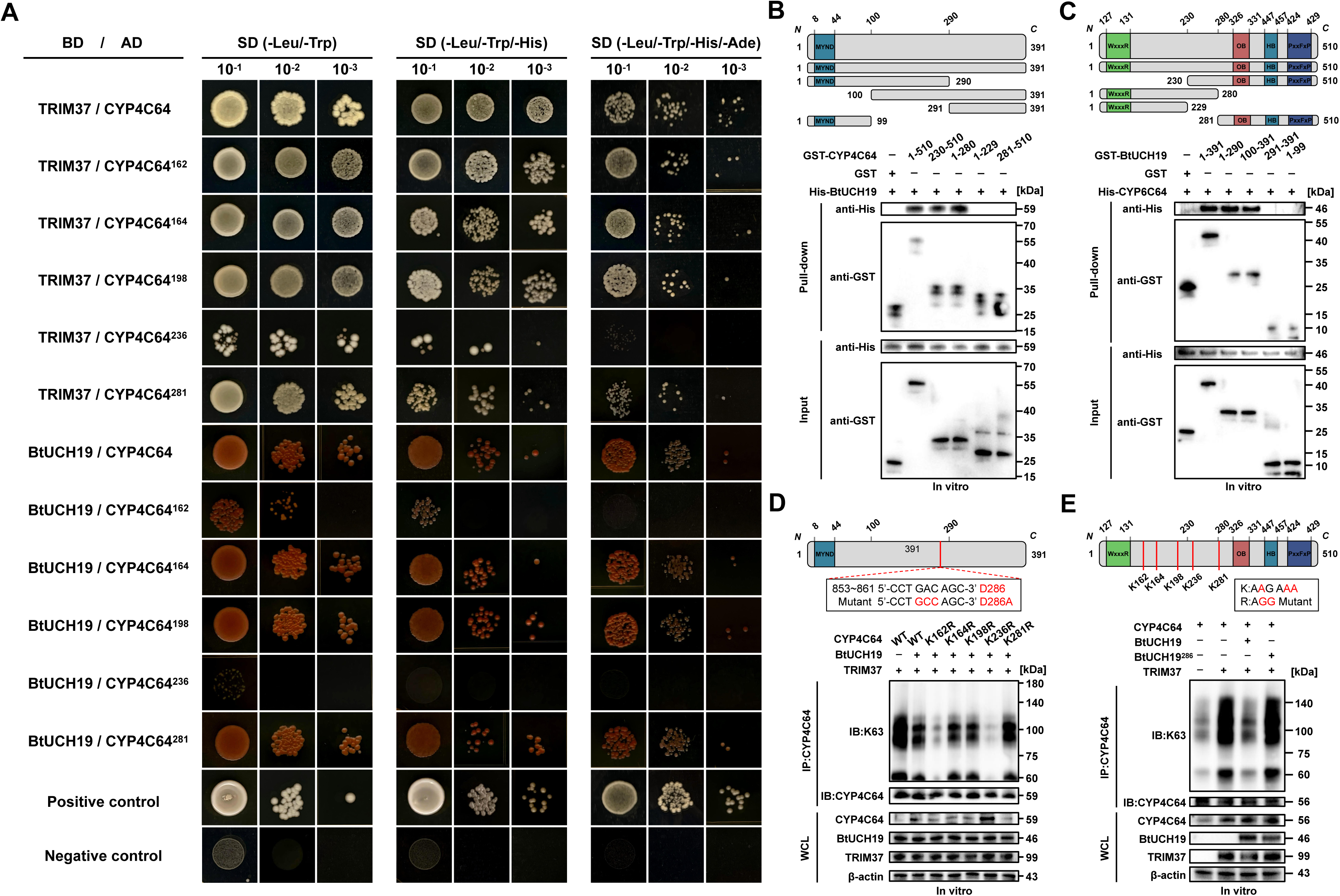
BtUCH19 promotes CYP4C64 deubiquitination through the K63 pathway. (A) Y2H assays showing the interaction between BtUCH19 / TRIM37 and CYP4C64 *in vitro*. CYP4C64 contains five mutations of potential ubiquitination sites, replacing the lysine (K) sites at positions 162, 164, 198, 236 and 281 of the N-terminal of CYP4C64 with arginine (R). SD, yeast drop-out culture medium. pGBKT7-53 and pGADT7-T were used as a positive control, pGBKT7-lam and pGADT7-T were used as a negative control. (B and C) Schematic diagram of CYP4C64 (B) and BtUCH19 (C) truncation mutation constructs (*above*). GST-Pull down analysis of the indicated proteins in BL21 transfected with the indicated plasmids of His-BtUCH19 and GST-CYP4C64 mutants (B) and His-CYP4C64 and GST-BtUCH19 mutants (C) (*Bottom*). (D and E) IP and western blotting analyses of the ubiquitination (K63) of CYP4C64 in S2 cells for 24 h; Schematic diagrams of CYP4C64 and BtUCH19 mutant constructs are shown (*Above*). The western blots tested the reciprocal interaction after mutation of CYP4C64 and BtUCH19 in transfected S2 cells, as detected by the signal intensity of CYP4C64 obtained with K48 antibody (*Bottom*). BtUCH19 interacts with CYP4C64 wild-type or CYP4C64 mutants (D); CYP4C64 interacts with BtUCH19 wild-type or BtUCH19 mutants (E). Note that AD (Activation domain), BD (DNA binding domain), SD (Synthetic dextrose minimal medium), - (lacking), Ade (Adenine), His (Histidine), Leu (Leucine), Trp (Tryptophan). WxxxR: WxxxR motif; OB: Oxygen-binding motif; HB: Heme-binding motif; PxxFxP: PxxFxP motif; MYND: Zinc finger domain. All western blot analyses represent at least three biological replicates and Input, *β*-actin and WCL (whole cell lysates) were used as loading controls. Cell lines in (D and E) were co-transfected to express TRIM37.

Based on *in silico* prediction using the AlphaFold 3 program (https://alphafold.com/) and PDBePISA (https://www.ebi.ac.uk/msd-srv/prot_int/), we performed simulated molecular docking of BtUCH19 and BtMYND619 with CYP4C64, respectively, and predicted the potential binding sites. It was observed that certain amino acid residues of CYP4C64 (Lysine / K-236) exhibited potential binding with specific amino acid residues of BtUCH19 (Asparagine / N-286) via the formation of hydrogen bonds and salt bridges (*SI Appendix*, Fig. S8A and Table. S10); Meanwhile, the binding sites between BtMYND619 and CYP4C64 are performed as a negative reference for docking (*SI Appendix*, Fig. S8B and Table. S10), BtMYND619 shows significantly worse than BtUCH19 in terms of interface area (area of protein interaction interface, Å), Δ^i^G (solvation free energy gain, Kcal/mol) and the number of chemical bonds (hydrogen bonds / N_HB_ and salt bridges / N_SB_), detailed information refer to *SI Appendix*, Table. S10.

To further explore the domains involved in the interaction between BtUCH19 and CYP4C64, we created truncation mutants of both proteins for use in GST-Pulldown assays (Fig. 6B, 6C and *SI Appendix*, Table. S7). The successful expression in *Escherichia coli* was confirmed by SDS-PAGE and Coomassie Brilliant Blue staining before co-incubation with the expressed protein (*SI Appendix*, Fig. S10A). The results revealed that BtUCH19 interacts with CYP4C64 containing the N-terminal residues 230-280, but does not interact with either the 1-229 fragment containing only the WxxxR motif (WxxxR) or the 281-510 fragment containing the oxygen-binding motif (OB), heme-binding motif (HB), and the conserved amino acid sequence PXXFXP near the carboxyl end (PxxFxP) (Fig. 6B). Similarly, upon truncating BtUCH19, it was found that CYP4C64 interacts with the N-terminal 100-290 region of BtUCH19, but does not interact with the 1-99 fragment containing only the zinc finger domain, nor with the 291-391 fragment of BtUCH19 (Fig. 6C). These results demonstrate that the N-terminal region of BtUCH19 (residues 100-290) and the N-terminal region of CYP4C64 (residues 230-280) are critical elements for their direct interaction and binding.

To further elucidate the site-specific role of BtUCH19 in regulating the function of CYP4C64, we leveraged the lysine to arginine mutant constructs above and transfected them into S2 cells for protein expression in combination with either TRIM37 or BtUCH19 (*SI Appendix*, Table. S7). Preliminary experiments showed that within 72 h after transfection of TRIM37 and CYP4C64 into S2 cells, CYP4C64 was ubiquitinated most highly at 12 h and was detected by antibodies K48 and K63 (*SI Appendix*, Fig. S10B). Coimmunoprecipitation experiments showed that the K162R and K236R mutations reduced BtUCH19-mediated CYP4C64 deubiquitination levels at 12 h after transfection (Fig. 6D and *SI Appendix*, Fig. S10C); In addition, the deubiquitination effect of BtUCH19 on CYP4C64 is dependent on its DUB enzyme activity, since a catalytically inactivated BtUCH19 mutant D286A is unable to deubiquitinate CYP4C64 in S2 cells (Fig. 6E and *SI Appendix*, Fig. S10D). Comparison of the data above with that generated for the K48 pathway (*SI Appendix*, Fig. S10C and S9D), demonstrates that the K63 pathway of CYP4C64 exhibits a stronger ubiquitination effect, indicating that TRIM37 and BtUCH19 enhance the ubiquitination and deubiquitination of CYP4C64 at K236. And this is also an important site for CYP4C64 to provide recognition for BtUCH19. Similarly, for BtUCH19, the core site for DUB enzyme function is D286 located in the N-terminal segment, and the degree of conservation of this site directly determines whether BtUCH19 can successfully deubiquitinate the K63 ubiquitin small molecule of the substrate. The above results clearly illustrate the molecular mechanism of the interaction between BtUCH19 and CYP4C64.

## Discussion

Our study unveils a novel "HGT-PTM-Metab" regulatory axis that drives insecticide resistance in *B. tabaci*, representing the first evidence of horizontal gene transfer (HGT) enabling xenobiotic adaptation through post-translational modification (PTM) in arthropods. Specifically, we demonstrate that the fungal-derived deubiquitinase BtUCH19-acquired via HGT-orchestrates insecticide resistance by stabilizing the key detoxification enzyme CYP4C64 through K63-linked deubiquitination. This PTM-mediated stabilization enhances the metabolic conversion of neonicotinoids into low-toxicity urea derivatives, establishing a direct mechanistic link from horizontally acquired regulatory genes to enhanced detoxification capacity (Fig. 7). This paradigm broadens the functional scope of HGT beyond terminal effector enzymes to include regulatory proteins of resistance pathways, while revealing a novel evolutionary path to insecticide resistance wherein horizontally transferred genes modulate endogenous resistance machinery. Our findings provide fundamental and applied insights into 1) horizontal gene transfer, 2) deubiquitination, and 3) P450-mediated insecticide metabolism. We discuss these topics below.

**Fig. 7.**
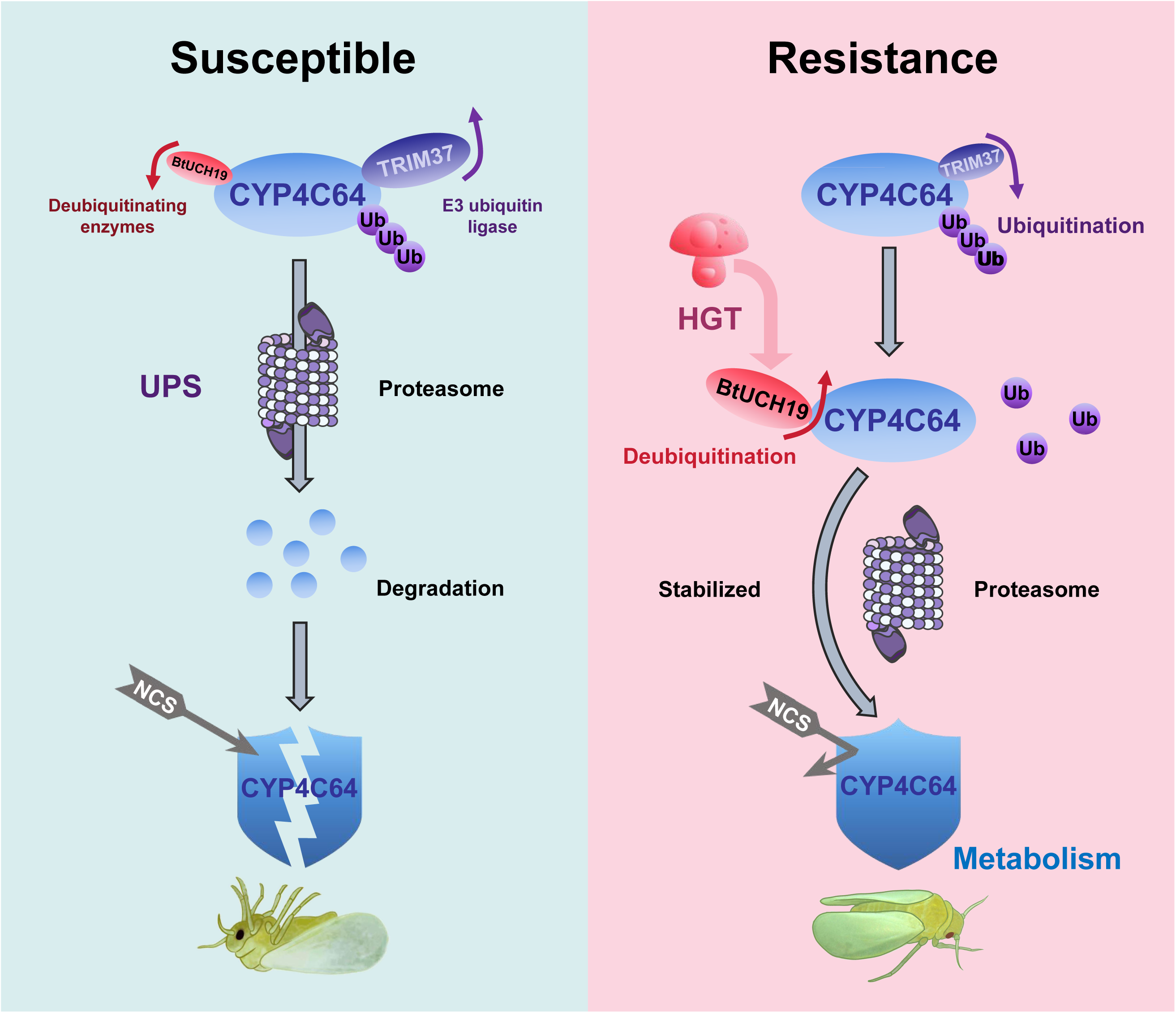
A horizontally transferred gene *BtUCH19* mediates CYP4C64 deubiquitination enhancing the resistance of *B. tabaci* to neonicotinoid insecticides. Upon horizontal transfer of a fungal gene *BtUCH19* to whiteflies, neonicotinoid resistant populations up-regulate the deubiquitinating enzyme gene *BtUCH19* and simultaneously down-regulate the E3 ubiquitin ligase *TRIM37*, which results in the deubiquitination of CYP4C64. This ensures sustained and stable high expression of a key detoxification enzyme *in vivo*, thereby enhancing resistance to neonicotinoid insecticides.

*B. tabaci* exhibits a very high frequency of HGT among insects (7), with fungal-derived HGTs playing prominent roles in nutrient acquisition (4, 5, 44, 45), immune defense (1, 46), and ecological adaptation (47). In *B. tabaci*, 78 HGT events originating from fungi have been identified (48, 49), primarily encoding direct-acting enzymes involved in feeding (50), detoxification (10), and immune defense against pathogens (51). Our functional characterization of a fungal-derived deubiquitinase, BtUCH19, reveals a novel mechanism of insecticide resistance (Fig. 2). Through K63-linked deubiquitination of the cytochrome P450 CYP4C64, BtUCH19 stabilizes this key detoxification enzyme, enhancing metabolic clearance of neonicotinoids. This represents the first demonstrated case of a HGT-derived factor involved in PTM regulation in arthropods, expanding the functional scope of horizontally acquired genes beyond direct metabolic catalysis to include modulation of endogenous resistance machinery. These findings also suggest that the role of HGT in other post-translational regulatory processes may be a fruitful area of future research.

Deubiquitination, a critical PTM, regulates protein stability and function by removing ubiquitin chains, influencing diverse biological processes. Although extensively studied in mammals, where deubiquitinating enzymes (DUBs) like USP6 and USP14 modulate signalling and proteasomal activity (11, 13, 52), its role in insects, particularly worldwide agricultural pests like *B. tabaci*, is less understood. Recent studies have highlighted the role of DUBs in immune signalling in the model insect *Drosophila*, with dUSP14 stabilizing immune regulators like Cactin by cleaving K63-linked ubiquitin chains (23). In *B. tabaci*, the genome encodes multiple DUBs, including USP, UCH, and OTU family members (Fig. S1B), some of which are upregulated under insecticide stress, suggesting a role in stress adaptation (53). We show that BtUCH19 acts as a key regulator of insecticide resistance by stabilizing the detoxification enzyme CYP4C64 through preferential K63 chain cleavage and K48 chain inhibition in the ubiquitin-proteasome system (UPS). BtUCH19 shares mechanistic parallels with known DUBs such as USP6 and USP14 in its ability to target K63-linked ubiquitin chains and modulate substrate fate (23, 52, 54). Like USP6 in JAK / STAT signaling or USP14 in proteasomal delay, BtUCH19 stabilizes and accumulates CYP4C64 under neonicotinoid stress conditions, which facilitates the metabolism of insecticides in whiteflies. However, compared to some endogenous whitefly deubiquitinating enzymes, BtUCH19 differs significantly in ubiquitin chain-type specificity, biological context, and evolutionary origin. The endogenous deubiquitinating enzymes in whiteflies are fewer and their functions are not strong enough. Taking *BtMYND619* as an example, regarding the capability of recognition to the key P450 detoxification enzyme CYP4C64, they are far inferior to the exogenous gene *BtUCH19*. This may be the endogenous driving force for whiteflies to select HGT.

These foreign genes significantly aid whiteflies in acquiring the ability to manipulate P450s, thereby enhancing their adaptability, especially in insecticide detoxification. The acquisition of *BtUCH19* via HGT thus not only introduces a new regulatory function but also fills a critical gap in the host’s ubiquitin-editing capacity.

The discovery of *BtUCH19* adds a new layer to our understanding of DUB-mediated regulatory networks in insects. Its K63 bias, upstream intervention role, and capacity to simultaneously suppress K63 and K48 ubiquitination on CYP4C64 support a finely tuned stabilization strategy under insecticide stress. Unlike generalist DUBs, BtUCH19 exemplifies functional precision acquired through HGT, revealing that detoxification regulation in pests may rely not only on metabolic enzymes but also on specialized regulatory factors, including detoxification enzymes (e.g., PMAT), protein-processing factors in whitefly (8–10, 48) and antibacterial genes in ladybugs (55). These findings highlight the importance of PTM-focused HGT events in shaping adaptive physiology and suggest that chain-specific DUBs such as BtUCH19 may represent viable targets for next-generation pest management strategies.

Cytochrome P450 monooxygenases (P450s) are critical detoxification enzymes in insects, with CYP4C64 playing a pivotal role in *B. tabaci* resistance to neonicotinoid insecticides (34, 56–59). Thiamethoxam (TMX) and clothianidin (CLO) are two widely used neonicotinoids, CLO is the *in vivo* activated form of TMX, and they share structural similarities, acting through the overactivation of nicotinic acetylcholine receptors (nAChRs) (41, 60–64), with CLO exhibiting a higher Toxic Equivalency Factor (Toxic Equivalency Factor / TEF ≈ 1.62) than TMX (TEF ≈ 0.11) due to its greater affinity for the nAChR (41, 65). The transformation of TMX to CLO represents a classical metabolic pathway through which TMX exerts its insecticidal effect in whiteflies, and similar ring-opening reactions have also been observed in insects, plants, and mammals (42, 66, 67). CYP4C64 counters this toxicity by metabolizing TMX and CLO into less toxic derivatives, a process enhanced by the deubiquitinase BtUCH19, highlighting a surprisingly sophisticated adaptation in whiteflies to counter insecticide stress.

We show that CYP4C64 converts TMX into CLO and thiamethoxam-urea (TMX-Urea), and independently metabolizes CLO into clothianidin-Urea (CLO-Urea), the derivatives of both ureas are final products which both exhibit significantly reduced toxicity to *B. tabaci* (40, 68–70). This enzymatic pathway begins with NADPH-CYP4C64-mediated cleavage of the aromatic heterocyclic ring of TMX, yielding CLO, with a higher nAChRs binding activity (71). This structural difference leads to CLO being accepted into the catalytic site of CYP4C64 with greater affinity than the parent TMX. Indeed, our kinetic analyses reveal that CYP4C64 exhibits a significantly higher V_max_ and lower K_m_ for CLO than TMX, resulting in a 2.7-fold greater K_cat_ (catalytic efficiency, Fig. 4B and *SI Appendix*, Table. S9), indicative of both enhanced turnover rate and stronger substrate affinity. This "toxicity-prioritized detoxification" strategy, as proposed by Nauen (72), allows whiteflies to quickly eliminate CLO.

The core role of CYP4C64 in this process is: With the assistance of the environment and a variety of enzymes (which may involve aldehyde oxidase, methyltransferase and nitroreductase, etc.)(66, 71), CYP4C64 disrupts the nitroguanidine group in TMX and CLO that possess nAChR toxic activity, reducing it to hydroxylamine group, and then catalyzes the demethylation oxidation reaction of the N-Methylguanidine group, causing hydrolysis of the C=N bond in the guanidino group, rearranging to form a carbonyl group, resulting in low-toxicity TMX-Urea and CLO-Urea (73, 74). These urea products are highly water-soluble, which greatly enhances the metabolic and excretion capacity of whiteflies to insecticides. We demonstrate that both NCS-Urea cause minimal interference with the insect’s nervous or metabolic systems and are excreted from the body via the honeydew of whiteflies, which follow a classic urea metabolism pathway in organisms (Fig. 4G) (40, 68, 75). Considering that the endogenous deubiquitinating enzymes of *B. tabaci* have weak recognition ability for the related P450s, the efficiency of this detoxification reaction is largely regulated by horizontally transferred genes, this special detoxification metabolic needs may be one of the driving forces behind HGT events.

Over a million years of evolution, herbivorous insects have developed sophisticated biological mechanisms to cope with the defensive chemicals produced by plants; these mechanisms have also enabled them to withstand the threats posed by human-applied pesticides (72, 76). The integration of HGT-derived regulatory mechanisms with an existing host-derived P450-mediated biotransformation system exemplifies synergistic evolution, enhancing whitefly resistance to neonicotinoids. By linking HGT, UPS regulation, and P450-mediated detoxification, our findings unveil a novel cross-kingdom regulatory mechanism underlying rapid insect adaptation (*SI Appendix*, Fig. S10). In conclusion, our data highlight the HGT-UPS as a key regulatory system that may enable insects to rapidly adapt to chemical stress, offering new insight into molecular strategies of resistance evolution. However, we should point out some limitations in this work, 1) What is the driving force for the horizontal transfer event of *BtUCH19* for whitefly? 2) Apart from the E3 ubiquitin ligase TRIM37, is CYP4C64 modified by other ubiquitinating enzymes (such as E1 and E2)? If so, what is the entire process of ubiquitination of CYP4C64? 3) Why does DUB-BtUCH19 exhibit a preference for constructing K63-type chains on substrates, over K48-type chains? To better understand why the whitefly acquired exogenous genes from fungi, including the impact of ubiquitination modification on the metabolic activity of CYP4C64, future research may need to compare BtUCH19 with other DUBs in whitefly systematically. In summary, studying PTMs and degradation pathways of enzymes that play a key role in pests, combined with tools like structural modeling, high-throughput screening, and substrate profiling, will enable the systematic mapping of DUBs functions in agricultural pests. Although pesticides are highly effective for pest control, neonicotinoid insecticides currently pose a significant threat to wild pollinators such as bees (77), this study will not only enhance our understanding of resistance driven by the UPS not only uncover new molecular targets for sustainable pest management, but also help us better protect vulnerable non-target beneficial insects.

## Materials and Methods

Detailed information on insect strains and cell lines, resistance level determination, q-PCR, RNAi experiments, Western blot, transgenic in *D. melanogaster*, heterologous expression, UPLC-MS, immunoprecipitation, RNAi-IP, Y2H, GST-Pull Down, and statistical analysis are described in *SI Appendix*, *SI Materials* and *Methods*.

## Supporting information

UCH19_Supplemental Information

## Acknowledgments

This research was supported by the National Natural Science Foundation of China (32361133558, 32221004, 32272598), China Agriculture Research System (CARS-24-C-02), Central Public-interest Scientific Institution Basal Research Fund (Y2023XK15; Y2024XK01). The Beijing Key Laboratory for Pest Control and Sustainable Cultivation of Vegetables and the Science and The Agricultural Science and Technology Innovation Program (ASTIP).

## Author contributions

C.B., X.Y., and Y.Z. designed research; H.L., C.Z., J.H., and C.W. performed research; R.Z., M.H., Q.T., C.Y., X.Z., and R.N. contributed new reagents/analytic tools; R.Z., M.H., Q.T., C.Y., and X.Y. analyzed data; and H.L., C.B., and X.Y. wrote the paper.

