## Supplementary material for "A horizontally transferred fungal deubiquitinase facilitates insecticides resistance in whitefly": UCH19_Supplemental Information

**This PDF file includes:**

Materials and methods
Figures S1 to S5
Tables S1 to S10
SI References

**Other supporting materials for this manuscript include the following:**

Datasets

### Supporting Information Text

#### Materials and methods

**Insects.** In order to collect the populations of *Bemisia. tabaci* suitable for the experiment, we tracked the dynamics of *B. tabaci* infestations across the country from 2021 to 2023 and conducted field sampling in vegetable-growing areas in China where only neonicotinoid insecticides are applied in large quantities. We collected 29 samples from field whitefly populations in nine provinces, with our laboratory susceptible strain a total of 30 populations. One lab-susceptible strain ($S^{\#1}$) has been maintained in the laboratory for more than 15 years without exposure to any insecticide. One field-susceptible population ($S^{\#2}$) and two field-resistant populations ($R^{\#1}$ and $R^{\#2}$) were identified based on the sensitivity to neonicotinoids in insecticide bioassays and were selected for the study. Details of the populations including information on the sampled location, sampling date, host plant and population identity are provided in *SI Appendix*, Table S1. The populations were identified by Zheng 's method (1) and were determined to be *B. tabaci* MED. To generate sufficient insects for subsequent tests, throughout the experiment, whiteflies were maintained in a suitable environment to simulate natural conditions for growth, development and reproduction. The rearing standards for populations of whiteflies were established according to relevant guidelines. Specifically, all whiteflies were kept in nylon cages located in a greenhouse with cotton (*Gossypium herbaceum* L. cv. Zhongmian 49) at an ambient temperature of 25±3°C, 70% relative humidity (RH) and a 14:10 light:dark (L:D) photoperiod. To prevent gene flow and exposure to insecticides, we placed nylon bags over the cage to isolate the susceptible populations. Resistant populations were maintained by watering cotton with a 2.0 mM aqueous solution of thiamethoxam every two months.

**Insecticides and Bioassays.** Insecticide bioassays were used to determine the sensitivity of *B. tabaci* populations to neonicotinoids and their metabolites (thiamethoxam, clothianidin, demethyl-thiamethoxam and clothianidin-urea).

Insecticides and metabolites were obtained from Shanghai Macklin Biochemical Co. To test the susceptibility of whiteflies to insecticides, bioassays were conducted on randomly captured adult males and females using the feeding method (2), with minor modifications. Specifically, for the diet solution, insecticides were dissolved in water to form a 10,000 mg / L aqueous solution, then the solution was diluted with raw diet solution (30.00% sucrose and 5.00% yeast extract, wt / vol) to serial dilutions of the tested concentrations (1.00-300.00 mg / L). The feeding device: Parafilm was used to cover the top of a glass tube (the tube length: 50 mm; outside diameter: 15 mm) and 40 μL of the diet solution was dropped onto the surface. Then, the second membrane was stretched over the first membrane, which created a chamber. Next, 30 healthy adult whiteflies were collected into tubes. Then the other opening of the tube was sealed with another Parafilm membrane. 40 μL of diet solution containing insecticides at different concentrations was set up, with four replicates per glass tube. For the control group, 40 μL of diet solution without insecticide was used. Finally, all tubes were placed in a black protective case and then placed in the environment above in a well-lit container. After 24 h / 48 h, the numbers of survivors and deaths were counted respectively and the mortality rate for each treatment was calculated.

**Resistance Level Determination.** To evaluate the resistance of *B. tabaci* to neonicotinoids, bioassays were performed on all samples. After establishing sensitivity baselines for susceptible populations, the resistance ratio (RR) was calculated by dividing the median lethal concentration ($LC_{50}$) of the susceptible lines by the $LC_{50}$ of the tested samples. The classification of resistance levels to insecticides was mainly defined according to Torres-Vila et al (3), with slight modifications:

RR<2 fold: susceptible; RR = 2-5 fold: minor; RR = 6-10 fold: low; RR =11-30 fold: moderate; RR = 31-160 fold: high; RR>160 fold: very high.

**Gene Cloning.** Total RNA was isolated from adult whiteflies following conventional TRIzol reagent methodology (Invitrogen, USA). The RNA purity and yield were quantified spectrophotometrically (NanoDrop 2000c, USA), after which genomic DNA-free RNA was reverse-transcribed into complementary DNA using oligo (dT) 18 primers and SuperScript II reverse transcriptase with integrated gDNA removal treatment (Takara, Japan). Gene sequences were computationally predicted through bioinformatic analysis of *B. tabaci* genomic databases and subsequently experimentally verified via real-time quantitative polymerase chain reaction (qPCR) amplification with sequence-specific primers (4), which are listed in the *SI Appendix*, Table. S4.

**Quantitative Real-time PCR (qPCR).** Quantitative real-time PCR (qPCR) was conducted on the Applied Biosystems 7,500 system with Power SYBR Green PCR Master Mix (Tiangen, China) in quadruplicate. Each 20 μL reaction mixture contained 10 μL 2× SuperReal PreMix Plus (Tiangen, China), 1 μL diluted cDNA, 0.6 μL each primer, 0.4 μL 50× ROX Reference Dye (Tiangen, China) and 7.4 μL dd$H_2O$. The thermal cycling protocol included an initial denaturation at 95°C for 1 min, followed by 40 cycles of 95°C for 5 s and 60°C for 15 s. Amplification efficiency was assessed using a twofold cDNA serial dilution standard curve. Gene expression fold-changes, normalized to housekeeping genes (*EF1α* and *RPL29* in *B. tabaci* and *RPL11* in *Drosophila melanogaster*), were calculated via the $2^{-\Delta\Delta CT}$ method (5). Three biological replicates (50 adults each per treatment) were analyzed. Primer sequences used are provided in the *SI Appendix*, Table. S4.

**Bioinformatic Analysis of Deubiquitinating Enzyme.** The *BtUCH19* gene was found in our sequenced *B. tabaci* MED genome (Gene ID: BTA013842.1, http://gigadb.org/dataset/100286) and annotated as a ubiquitin Carboxyl-Terminal Hydrolase Isozyme. For the cloning of *BtUCH19*, the putative CDS of *BtUCH19* was manually corrected using our previously completed transcriptome data of *B. tabaci* MED (6). Specific primers used for gene cloning (*SI Appendix*, Table. S4) were designed using Primer Premier 5.0 and the PCR amplicon from *B. tabaci* MED was cloned into the pEASY-T1 (TransGen, China) vector and transformed into *Escherichia coli* Trans1-T1 competent cells (TransGen, China) for sequencing.

The full-length cDNA sequence of *BtUCH19* was deposited in the GenBank database (accession number XM_072302559).

To confirm that *BtUCH19* is indeed a horizontally transferred gene, we conducted the following analysis. Following gene cloning, conserved protein domains were predicted using the NCBI CDD database. Transmembrane domains (TMDs) were identified using TMHMM, while theoretical isoelectric points (pI) and molecular weights (Mw) were calculated using ExPASy ProtParam. Homologous sequences (XM_072302559.1) and upstream (XP_018899910.1) / downstream (XM_072302585.1) sequences retrieved from NCBI were aligned and used to construct a phylogenetic tree in MEGA11 (https://www.megasoftware.net/) using the neighbor-joining method with 10,000 bootstrap replicates *Supplementary Material*, Fig. S1B and S1C. To identify homologous sequences of *BtUCH19*, PSI-BLAST (Position-Specific Iterated BLAST 2.15.0+) searches were performed against a locally constructed non-redundant (nr) protein database using the *BtUCH19* amino acid sequence as the initial query. The initial BLAST search was limited to the top 100 hits based on E-value and sequence similarity. These sequences were then used to build a position-specific scoring matrix (PSSM), which was employed in two additional PSI-BLAST iterations, resulting in a total of three rounds of searching to enhance sensitivity and retrieve remote homologs. The resulting sequences were aligned using MAFFT (v7.520) with default parameters. Global alignment (globalxx) and local alignment (localms) are used to calculate the percent identity between each reference sequence and the target sequence. The sequence-related information used for constructing the phylogenetic tree is listed in *Supplementary Material*, Fig.S1. A phylogenetic tree was constructed using FastTree (version 2.1.11, No SSE3), which applies the approximately-maximum-likelihood method for large alignments. The generated tree was visualized and annotated using the Interactive Tree of Life (iTOL) online tool (https://itol.embl.de/). The endogenous deubiquitinating gene *BtMYND619* of whiteflies was used as the control gene, and the same analysis was conducted with reference to BtUCH19.

**Western Blot.** Protein extraction was performed on 300 adult whiteflies per sample using the ProteinExt Total Protein Extraction Kit (Beyotime, China). Quantified protein lysates (BCA assay, Beyotime, China) were denatured with SDS-PAGE loading buffer (CWBIO, China) at 100°C for 10 min, separated on 4-20% gradient precast gels (Lablead Biotech., China) and transferred to PVDF membranes (Merck Millipore, Germany). Membranes were probed overnight at 4°C with rabbit polyclonal primary antibodies against target proteins (*SI Appendix*, Table. S8), followed by HRP-conjugated goat anti-rabbit secondary antibody (1:5K, CWBIO, China). Chemiluminescent signals were detected using SuperSignal West Pico substrate (Thermo Fisher, USA) and imaged on a Tanon-5200 system. Primary antibodies included: CYP4C64 (1:1.5k), BtUCH19 (1:1.5k), TRIM37 (1:5k). Antibodies were either custom-generated against synthetic peptides or commercially sourced, with specificity validated by peptide-BLAST against the whitefly genome. *β*-actin (1:5k, Abcam, USA) served as the loading control.

**RNAi Experiments.** Yang et al. 's method was used with minor modifications (2). Double-stranded RNAs (dsRNAs) were synthesized *in vitro* using the T7 RiboMAX™ Express RNAi System (Promega, USA) following standardized protocols. Gene-specific primers (*SI Appendix*, Table. S4) containing T7 promoter sequences were employed for template amplification before *in vitro* transcription. Newly emerged adult *B. tabaci* were subjected to oral administration of dsRNAs at 24 h post-eclosion. Post-treatment, specimens were selectively collected for molecular analyses, including total RNA extraction using TRIzol reagent and cDNA synthesis with reverse transcriptase. qPCR was performed as detailed above using SYBR Green chemistry to assess RNA interference (RNAi) efficiency through quantification of target gene expression levels relative to endogenous controls. dsRNA treatments demonstrating statistically significant knockdown ($p<0.05$) were subsequently utilized for phenotypic characterization. All experiments included parallel negative controls treated with dsRNA targeting the non-homologous sequences *EGFP* (enhanced green fluorescent protein).

**Transgenic Expression of the *CYP4C64* Gene in *D. melanogaster*.** The wild-type *CYP4C64* gene was synthesized and cloned into the *pUASTattB40* plasmid (Thermo Fisher, USA). Germline transformation of *Drosophila melanogaster* embryos carrying the *attP40* docking site on chromosome 2 [genotype: $w^{(1118)}$*; P{w(+mC) = UAS-CYP4C64}attP40(25C6)/CyO*] was performed via PhiC31 integrase-mediated site-specific recombination (BestGene Inc., USA). Transgenic lines were genetically stabilized using balancer chromosomes and successful integration of *CYP4C64* was verified through PCR amplification with LA Taq DNA polymerase (TaKaRa Bio, Japan), followed by Sanger sequencing using primers listed in *SI Appendix*, Table. S4. To drive tissue-specific expression, virgin female *Tub-GAL4* driver flies [genotype: $w^{(1118)}$*; P{w(+mC) = act5C-gal4}attP2 (68A4)/TM6B*] were crossed with male *UAS-CYP4C64* transgenic flies (FunGene Biological, China). For toxicological assessment, F1 adults (2-5 days post-eclosion) were exposed to neonicotinoids (Sigma-Aldrich) dissolved in aqueous 1% sucrose solution layered onto 2% agar in standard *Drosophila* vials. Insecticide concentrations were air-dried overnight at 25°C under sterile conditions. Cohorts of 10-20 flies per vial were subjected to four biological replicates per concentration, with mortality quantified 48 h post-exposure using a standardized scoring protocol. All bioassays were conducted under controlled environmental conditions (25 ± 1°C, 60% RH, 12:12 / L:D photoperiod) to minimize confounding variables. Using vials containing agar / sucrose without insecticide as control mortality.

**Heterologous Expression of *CYP4C64* and Activity Measurements.** The Heterologous Expression of CYP4C64 and BtCPR was assessed by the previous method (Lu *et al.*). *B. tabaci CYP4C64* and *B. tabaci* cytochrome P450 reductase (*BtCPR*) were PCR amplified using the primers and Q5.High-Fidelity DNA Polymerase (TIANGEN, China) as described above (*SI Appendix*, Table. S4). The Bac-to-Bac baculovirus expression system was utilized to functionally express CYP4C64 and BtCPR *in vitro* by cloning the gene into the pFast-Bac HTA vector via the Clon Express Ultra One-Step Cloning Kit (Vazyme, China) and transferring the recombinant plasmid into DH10Bac competent cells for bacmid construction

(Solarbio, China, *SI Appendix*, Table. S7). Purifying endotoxin-free bacmid DNA (pJFRC28-10XUAS-IVS-p10, *SI Appendix*, Table. S7) using the EndoFree Maxi Plasmid Kit (TIANGEN, China). Transfecting Spodoptera frugiperda clone 9 (Sf9) cells with FUGENE reagent to generate recombinant baculovirus particles (P1, P2 and P3) (Promega, USA). The $P3^{CYP4C64}$ was co-expressed with $P3^{BtCPR}$ in High Five / BTI-Tn-5B1-4 cells (H5), then harvested 44 h post-transfection and washed with 0.1 M PBS (Solarbio, China). Then P450 protein-microsomes were prepared as described previously, detecting P450s activity using the reduced CO-difference spectra of recombinant test, the concentration of P450 protein was calculated by referring to the following formula (7):

$$\frac{A_{450} - A_{490}}{0.091^{*}} \times \text{Dilution ratio} = {}^{\text{nmol P450}}\!/_{\text{mg protein}}$$

$^{*}$ The molar extinction coefficient of P450 ($\mu M - 1cm^{-1}$)

The proteins extracted from cells mock-transfected were used as a control group.

**P450 Catalytic Assays.** P450s *in vitro* metabolism was assessed by the previous method (Lu *et al.*). The concentration of 50 pmol recombinant P450 proteins was prepared, with an NADPH regeneration system (Promega, USA) and 1 μL neonicotinoid standard (0.1 mg / mL, *SI Appendix*, Table. S11) constituting the metabolism reaction. The reaction was incubated at 30°C in a shaking table at 1200 rpm / min for 1 h. Thereafter, metabolic reaction products were quenched by centrifugation (20,000 ×g, 10 min) and supernatants filtered through 0.22 μm nylon membranes (Jinteng, China) for UPLC-MS analysis. P450s Microsomes incubated without NADPH were used as a control. The data was fit to the Michaelis-Menten equation using non-linear regression to calculate related enzyme kinetic constants such as $V_{max}$ (maximum rate of metabolism), $K_m$ (Michaelis constant) and $K_{cat}$ (Catalytic constant: $V_{max}$ / $K_m$).

**UPLC-MS Analysis of Neonicotinoids.** The Ultra-performance liquid chromatography-mass spectrometry (UPLC-MS) is used for the analysis of neonicotinoids and their metabolites. Metabolic reaction products were resolved on an Atlantis T3 column (2.1 × 50 mm, 3 μm; Waters, USA) using a Waters ACQUITY UPLC I-Class / Xevo TQ-S Micro system. Mobile phases, gradient elution were assessed by previous condition (Lu *et al.*), Column and autosampler temperatures were maintained at 30°C and 4°C, respectively. Electrospray ionization in positive mode (ESI+) with MRM transitions was used to quantify neonicotinoids. Data were processed using Progenesis QI (v3.0) with photodiode array detection (190-800 nm, mass accuracy < 5 ppm).

To elucidate the specific parameters of CYP4C64-mediated neonicotinoids and their metabolites, we set a series of time gradients (0-60 min) and neonicotinoid concentration gradients (200-1 pmol) and detected the peak area change of the neonicotinoids and their metabolites in each reaction. The incubation time for the concentration gradient experiment was selected based on the time point with the highest metabolic efficiency from the time gradient experiment.

To identify the metabolites of neonicotinoids, we selected commonly available insecticide metabolites and conducted UPLC-MS analysis. We then compared the product peaks observed after CYP4C64-mediated metabolism of the neonicotinoid substrates in this experiment with those obtained from the purchased standards of metabolites to determine their specific types. Quantification was performed by comparison with the standard curve obtained from pure running standards of metabolites as part of the same batch of samples. The ratio of metabolites to neonicotinoids was calculated based on the standard curves. Detailed information and monitoring parameters on substrates and their metabolites are listed in *SI Appendix*, Table. S1 and S2. Microsomes incubated without NADPH were used as the control group.

**Honeydew analysis.** To further investigate whether *CYP4C64* / *BtUCH19* / *TRIM37* / *BtMYND619* are responsible for the metabolism of neonicotinoids in *B. tabaci*, we conducted RNA interference to relative genes. Following this, honeydew was collected and analyzed via UPLC-MS after feeding whiteflies TMX and CLO for 48h. Two insecticides are dissolved simultaneously in the diet solution at a final concentration of 0.5 mM together. The honeydew collection method refers to Xia's method (8). Specifically, the honeydew collection device consists of a vertically oriented glass tube and a plastic bottom covered with tinfoil and each tube includes 10 male and 40 female. The collected honeydew was dissolved in 500 μL of 80% methanol aqueous solution to reach a final concentration of 0.1 g/mL. The supernatants were then filtered through 0.22 μm nylon membranes (Jinteng, China) for UPLC-MS analysis. Whiteflies fed on ds*EGFP* serving as a control and quantitative calculations were performed by referring to the standard curves, each test representing at least six replicates.

**S2 Cell Culture and Transfection.** *Drosophila melanogaster* S2 embryonic cells (S2) were cultured in Hyclone SFX-Insect Medium (Thermo Fisher, USA) under aseptic conditions at 27°C. Routine subculturing was performed every 48 h by mechanical detachment and resuspension in fresh medium to maintain logarithmic growth. For transfection experiments, cells were seeded into 6-well plates at a density of $6 \times 10^5$ cells per well in 300 μL of pre-equilibrated medium and allowed to adhere for 1 h before transfection. Lipofectamine 2000 (Invitrogen, USA) was utilized for plasmid delivery following optimized protocols. Plasmid DNA (100 ng / μL in Tris-EDTA buffer) and transfection reagent were mixed at a volumetric ratio of 1:3 (6 μL plasmid:2 μL lipofectamine 2000) in 100 μL of serum-free medium, followed by 20 min incubation at ambient temperature to form transfection complexes. The DNA-lipid complexes were then gently overlaid onto adherent S2 cells. Post-transfection, plates were maintained at 27°C in a humidified incubator for 48 h without medium replacement to ensure sustained gene expression. Transfection efficiency was monitored via parallel wells transfected with a GFP-expressing control plasmid. All experimental conditions included triplicate wells and were normalized against mock-transfected controls containing lipofectamine 2000 alone.

**Immunoprecipitation (IP) Assays.** Immunoprecipitation assays were performed to assess direct protein interactions *in vivo* using total protein lysates from the insecticide-resistant *B. tabaci* population $R^{\#2}$. Proteins were extracted with the Total Protein Extraction Kit (Beyotime, China) in the presence of protease inhibitors (1×Protease Inhibitor, Beyotime, China) (1:100 dilution) to maintain protein integrity. For immunoprecipitation, pre-cleared lysates (500 μg per reaction) were incubated overnight at 4°C with 5 μg of primary antibodies targeting protein, followed by a 3 h incubation with Protein A / G magnetic beads (BEAVER, China) to capture immune complexes. The bead-bound complexes were sequentially washed under stringent conditions: three washes with IP lysis buffer (50 mM Tris-HCl pH 7.4, 150 mM NaCl, 1% NP-40), two washes with high-salt buffer (IP buffer supplemented with 500 mM NaCl) and a final rinse with PBS (pH 7.4) to eliminate nonspecific interactions. Bound proteins were eluted through three cycles of 10 min incubation with 0.1 M glycine-HCl (pH 2.5) under vigorous agitation, followed by centrifugation (12,000 ×g, 4 °C, 5 min) to collect eluates. Neutralization was achieved by adding 1 M Tris-HCl (pH 8.0) at a 1:10 ratio to stabilize eluted proteins. The immunoprecipitation samples were separated by 10% SDS-PAGE followed by western blot analysis to detect the precipitation of target proteins. The IgG antibody was used as a negative control.

**RNAi-IP Assay.** To investigate the ubiquitination and deubiquitination of CYP4C64 by TRIM37 and BtUCH19, adult individuals of the resistant *B. tabaci* population $R^{\#2}$ were subjected to RNAi by oral delivery of dsRNA targeting *TRIM37* and *BtUCH19* transcripts. Following dsRNA administration, whole-body protein lysates were extracted under denaturing conditions and subjected to immunoprecipitation (IP) using anti-CYP4C64 antibodies, as previously described. Immunoprecipitated proteins were resolved by SDS-PAGE and transferred to PVDF membranes for western blot analysis. CYP4C64 was detected using a rabbit polyclonal anti-CYP4C64 primary antibody (1:1.5K).

**Enzyme Activity Assays.** BtUCH19 deubiquitinase activity was measured according to the protocol of the Deubiquitinase Assay Kit (Abcam, UK). For DUB protease assays referred to V. Vozandychova et al. 's method (9), whitefly proteins were extracted with the Total Protein Extraction Kit (Beyotime, China) in the presence of protease inhibitors (1×Protease Inhibitor, Beyotime, China) (1:100 dilution) to maintain protein integrity. Then collected protein was lysed in cold assay buffer containing 1 mM DTT, centrifuged at 10,000×g at 4°C for 15 minutes and the supernatant collected. The supernatant was then immunoprecipitated (IP) with anti-BtUCH19 antibody and eluted overnight at 4°C, collecting the acquisition of BtUCH19 protease with deubiquitinating activity. Following the addition of the substrate mixture, fluorescence intensity was measured using the microplate provided in the kit in conjunction with a microplate reader (Bio-Rad, USA). The measurements were taken at 350 nm excitation and 440 nm emission to analyze activity. PBS was used as no-substrate control.

To examine the impact of knockdown of *BtUCH19* on enzyme activity, ds*BtUCH19* was used for RNAi feeding of 300 adult whiteflies. Samples were collected 6 hours later and rapidly frozen with liquid nitrogen for homogenization. Subsequently, the total protein was extracted and used to assess enzyme activity as follows: Cytochrome P450 enzyme activity was measured according to the protocol of the P450s Assay Kit and referencing Wang et al. 's method (10) (Ziker, China). Glutathione S-transferase / GST was assessed using GST ELISA assay kit (Solarbio, China) and UDP-glycosyltransferase / UGT activity was assessed using the Insect UGT ELISA assay kit (LMAI BIO, China) to measure the detoxification enzyme activity in whole *B. tabaci* adults according to the manufacturer's instructions (11, 12). Insects fed with ds*EGFP* served as the control group.

**Yeast Two-Hybrid (Y2H) Assays.** Yeast two-hybrid assays were conducted to investigate protein-protein interactions *in vitro*. The coding sequences (CDS) of *CYP4C64* and *CYP4C64* mutants were subcloned into the pGADT7 activation domain (AD) vector to generate AD-*CYP4C64* and AD-*CYP4C64* mutant prey constructs, while the CDS of *TRIM37* and *BtUCH19* were ligated into the pGBKT7 DNA-binding domain (BD) vector to produce BD-*TRIM37* and BD-*BtUCH19* bait constructs (*SI Appendix*, Table. S7). All bait-prey plasmid combinations were co-transformed into *Saccharomyces cerevisiae* strain Y2HGold (Clontech, Japan) via the lithium acetate method. Transformed yeast cells were serially diluted (1:10, 1:100 and 1:1000) in sterile PBS and spotted onto three different selection synthetic dextrose medium (SD, lacking Adenine, Histidine, Leucine and Tryptophan) to assess interaction-dependent growth. Images were captured on day 5 following incubation at 30°C. Positive controls (pGBKT7-53 + pGADT7-T) and negative controls (pGBKT7-lam + pGADT7-T) were included in parallel to validate system specificity. Protein interactions were confirmed by observable yeast growth under selective conditions, with interaction strength correlated to colony density across a dilution gradient.

**GST-Pull Down.** *BtUCH19*, *CYP4C64* and their truncated segments were separately subcloned into the pGEX-4T and pet28a vectors (*SI Appendix*, Table. S7). The recombinant vectors were subsequently transformed into the *Escherichia coli* strain BL21 and protein expression was subsequently induced for 12 hours with 0.5 mM isopropyl-β-d-thiogalactopyranoside (Invitrogen, USA) at 15°C, 120 rpm to produce the activated GST-CYP4C64, GST-BtUCH19, His-CYP4C64, His-BtUCH19 and their labeled truncated fragment recombinant proteins. The empty pGEX-4T and empty pet28-a vectors were used to produce the GST and His proteins, respectively. The effect of IPTG-induced protein expression was identified through SDS-PAGE combined with Coomassie Brilliant Blue staining. The GST and GST fusion proteins were purified using glutathione-conjugated agarose beads (Thermo Fisher, USA).

His fusion proteins were purified using nickel-nitrilotriacetic acid (NiNTA) agarose beads (Thermo Fisher, USA). Glutathione agarose beads (50 μL) coated with either GST-*CYP4C64* or GST-*BtUCH19* recombinant protein (5 mg per sample) were mixed with the His-tagged recombinant proteins (10 mg per sample) separately in 500 μL of PBS (pH 8.0). Each mixture was incubated with shaking for 6 hours at 4°C. The beads were eluted with 200 μL of 50 mM tris-HCl (pH 8.0) containing 10 mM GSH to isolate the supernatant following three washes. Detailed information about the prokaryotic expressed protein *in vitro* is listed in (*SI Appendix*, Table. S7), and the eluted samples were subjected to western blot analysis with Anti-His (1:5k) and Anti-GST (1:5k).

**Modeling of Protein Interaction.** We predicted BtUCH19 / BtMYND619 and CYP4C64 interactions using computational tools. Briefly, protein models were created using the online AlphaFold 3 (https://alphafoldserver.com/) program and the coupled amino acid sequences of BtUCH19 / BtMYND619 and CYP4C64. The PDBePISA program (https://www.ebi.ac.uk/msd-srv/prot_int/pistart.html) was used for predicting docking interactions. The results were filtered on the basis of the calculation of the docking scores using ITScorePP to obtain the most reliable docking models. PyMOL software (version 3.0.3) was used for analysis and drawing operations.

**Statistical Analysis and Data Visualization.** The raw data were analyzed as follows: $LC_{50}$ values were calculated from the bioassay data and their 95% confidence limits according to probit regression via POLO PLUS 2.0 software (LeOra Software, USA). The statistical analyses were performed with SPSS 26.0.0.0 (SPSS, USA). The analysis of variance (ANOVA) with Tukey's post hoc test (GraphPad 8.0). $K_m$, $V_{max}$ and $K_{cat}$ were determined using GraphPad 7.0 (Insightful Science, USA) by fitting data to the Michaelis-Menten equation via non-linear regression. All quantitative data are reported as means ± SEM from at least three independent experiments.

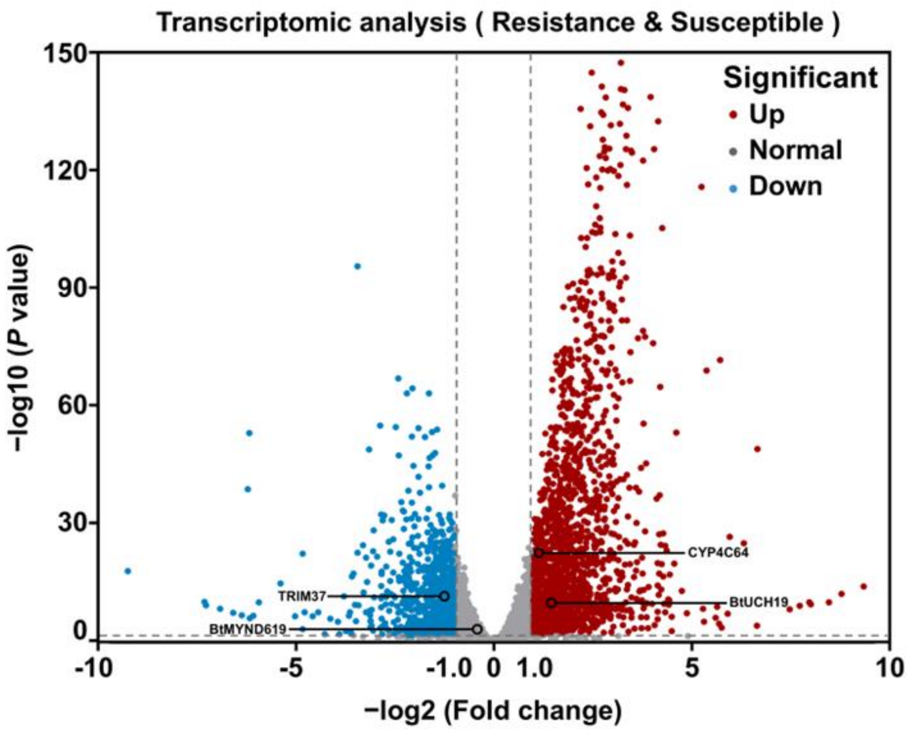


**Fig. S1. *BtUCH19* is upregulated in the genome of *B. tabaci* MED.** Transcriptomic analysis of *BtUCH19* expression in resistant and susceptible populations of *B. tabaci* MED. Note that the plotted data were mined from the Yang Li's database (Y. Li *et al*. 2022) and the transcriptome database related to *B. tabaci* MED insecticide resistance (http://www.whiteflygenomics.org/ftp/MED/). The stars indicate *BtUCH19* of the whitefly *B. tabaci* MED, MED: Mediterranean.

Note that the relevant data has been uploaded to NCBI (BioProject: PRJNA1401507)

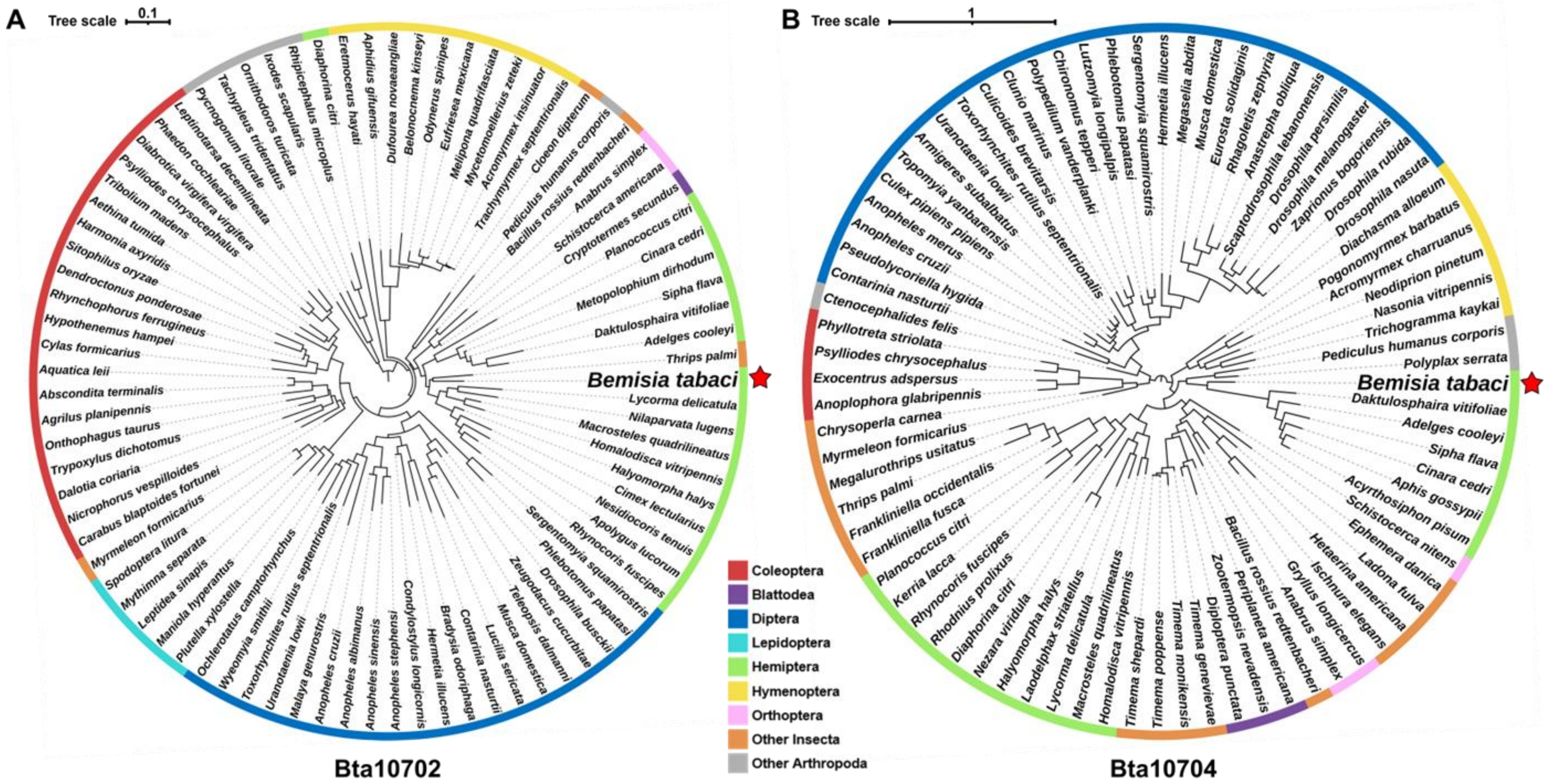


**Fig. S2. Phylogenetic analysis of the upstream and downstream genes of BtUCH19 from arthropods.** (A) *Bta10702* is the upstream (5') gene of *BtUCH19*, was identified as a fasciculation and elongation protein zeta-2 isoform X2 gene (450 bp, Gene Bank: XP_072158686.1). (B) *Bta10704* is the downstream (3') gene of *BtUCH19*, was identified as a very-long-chain (3R)-3-hydroxyacyl-CoA dehydratase hpo-8 gene (Gene Bank: XP_018899910.1). The order to which different species belong is indicated by different colors. The red stars indicate genes of the whitefly *B. tabaci*

MED (Mediterranean). *BtUCH19* is flanked by two typical arthropod genes, demonstrating the specificity of its origin. Note that all phylogenetic trees were constructed in MEGA11 (https://www.megasoftware.net/) from MAFFT-aligned sequences retrieved by PSI-BLAST, using FastTree with the maximum likelihood method, then visualized via iTOL (https://itol.embl.de/).

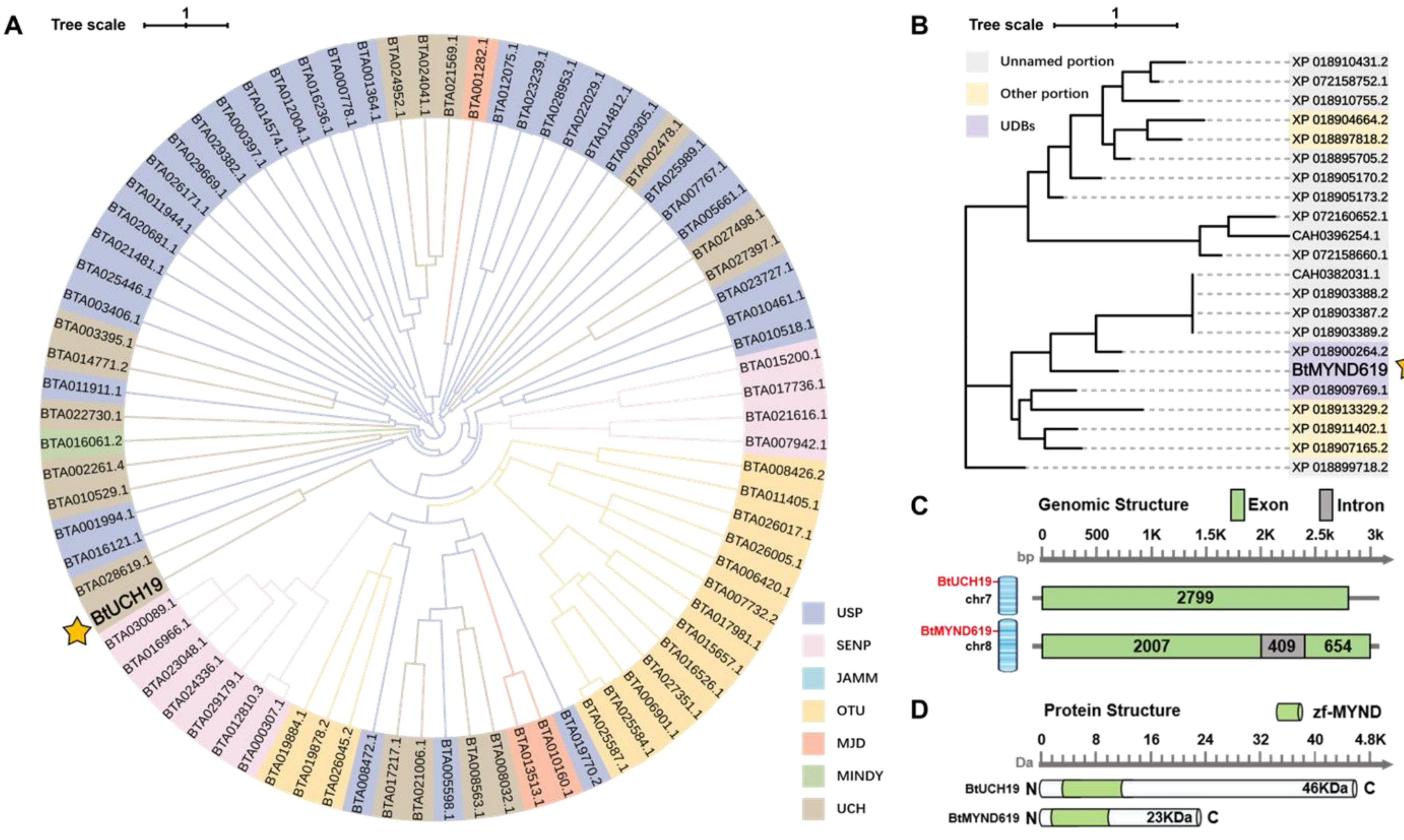

A
Tree scale
1
BTA024952.1
BTA024041.1
BTA021569.1
BTA001282.1
BTA012075.1
BTA023239.1
BTA028953.1
BTA022029.1
BTA014812.1
BTA009305.1
BTA002478.1
BTA025989.1
BTA007767.1
BTA005661.1
BTA027498.1
BTA027397.1
BTA023727.1
BTA010461.1
BTA010518.1
BTA015200.1
BTA017736.1
BTA021616.1
BTA007942.1
BTA008426.2
BTA011405.1
BTA026017.1
BTA026005.1
BTA006420.1
BTA007732.2
BTA017981.1
BTA015657.1
BTA016526.1
BTA027351.1
BTA006901.1
BTA025584.1
BTA025587.1
BTA019770.2
BTA010160.1
BTA013513.1
BTA008032.1
BTA008563.1
BTA005598.1
BTA021006.1
BTA017217.1
BTA008472.1
BTA026045.2
BTA019878.2
BTA019884.1
BTA000307.1
BTA012810.3
BTA029179.1
BTA024336.1
BTA023048.1
BTA016966.1
BTA030089.1
BtUCH19
BTA028619.1
BTA016121.1
BTA001994.1
BTA010529.1
BTA002261.4
BTA016061.2
BTA022730.1
BTA011911.1
BTA014771.2
BTA003395.1
BTA003406.1
BTA025446.1
BTA021481.1
BTA020681.1
BTA011944.1
BTA026171.1
BTA029669.1
BTA029382.1
BTA000397.1
BTA014574.1
BTA012004.1
BTA016236.1
BTA000778.1
BTA001364.1
USP
SENP
JAMM
OTU
MJD
MINDY
UCH
B
Tree scale
1
Unnamed portion
Other portion
UDBs
XP 018910431.2
XP 072158752.1
XP 018910755.2
XP 018904664.2
XP 018897818.2
XP 018895705.2
XP 018905170.2
XP 018905173.2
XP 072160652.1
CAH0396254.1
XP 072158660.1
CAH0382031.1
XP 018903388.2
XP 018903387.2
XP 018903389.2
XP 018900264.2
BtMYND619
XP 018909769.1
XP 018913329.2
XP 018911402.1
XP 018907165.2
XP 018899718.2
C
Genomic Structure
Exon
Intron
0
500
1K
1.5K
2K
2.5k
3k
bp
BtUCH19
chr7
2799
BtMYND619
chr8
2007
409
654
D
Protein Structure
zf-MYND
0
8
16
24
32
40
4.8K
Da
BtUCH19 N
46KDa
C
BtMYND619 N
23KDa
C

**Fig. S3. Bioinformatic analysis of deubiquitinating enzymes in whitefly.** Analyses of the *BtUCH19* and *BtMYND619* in *B. tabaci* MED. (A) Phylogenetic tree demonstrates that *BtUCH19* belongs to the ubiquitin carboxyl-terminal hydrolase family in *B. tabaci* MED. *BtMYND619* is the deubiquitinating enzyme with the highest similarity to *BtUCH19*. (B) Phylogenetic tree demonstrates that fewer deubiquitinating enzymes similar to BtMYND619 are present in *B. tabaci* MED. (C and D) Position and domain composition of *BtUCH19* and *BtMYND619* in *B. tabaci* MED. BtUCH19 is located on chromosome 7 of the whitefly genome, with a length of 2799 bp and contains one exon, and *BtMYND619* is located on chromosome 8 of the whitefly genome, with a length of 2985 bp and contains two exons and one intron (C). There is a zinc finger domain (zf-MYND), which is commonly found in deubiquitinating enzymes at the N-terminus of both *BtUCH19* and *BtMYND619* (D). Note that all phylogenetic trees were constructed in MEGA11 (https://www.megasoftware.net/) from MAFFT-aligned sequences retrieved by PSI-BLAST, using FastTree with the maximum likelihood method, then visualized via iTOL (https://itol.embl.de/). MED: Mediterranean.

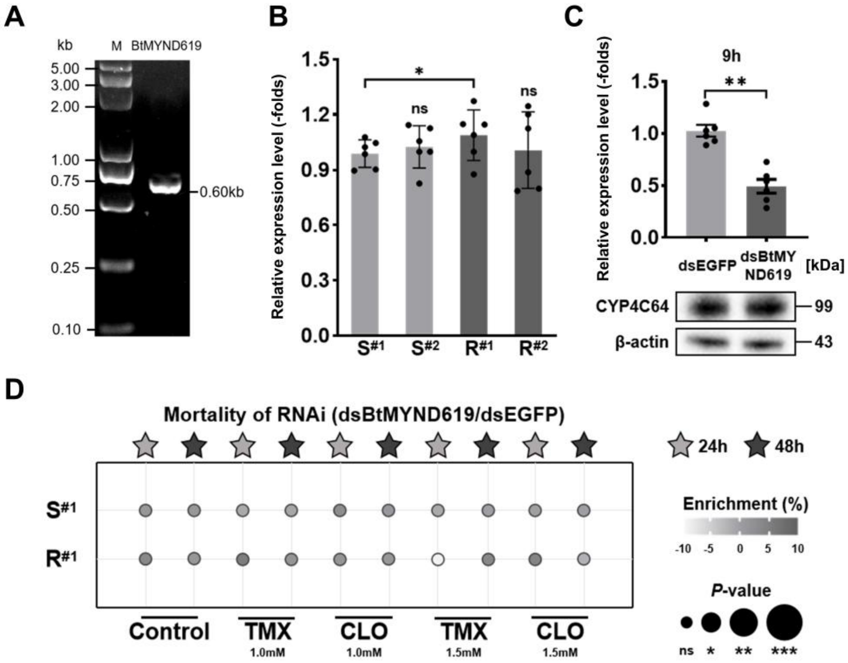


**Fig. S4. *BtMYND619* does not participate in the abundance of CYP4C64.** (A) The *BtMYND619* was detected using an SDS-PAGE precast gel. (B) Quantification of the expression of the mRNA of *BtMYND619* among the experimental populations using qPCR analyse. (C) Relative expression analyses of *BtMYND619* in mRNA level and CYP4C64 in protein level after feeding on ds*BtMYND619* for 9 h in $R^{\#1}$. (D) Sensitivity of the experimental populations to TMX and CLO after RNAi knockdown of *BtMYND619*. Note that all qPCR data are presented as the mean ± SEM of at least three independent experiments. All bioassay data (mean ± SEM) represent at least three biological replicates with approximately 60 adults in each replication and mortality was assessed 24 h and 48 h after first exposure to insecticide, and newly emerged (1-day-old) adults were used in the RNAi experiments and whiteflies fed on ds*EGFP* were used as a control. All western blot analyses represent at least three biological replicates and *β*-actin was used as a loading control. Data were analyzed with ANOVA with Tukey's HSD post hoc test: ns (not significant) $P > 0.05$, $^{*}P < 0.05$ and $^{**}P < 0.01$.

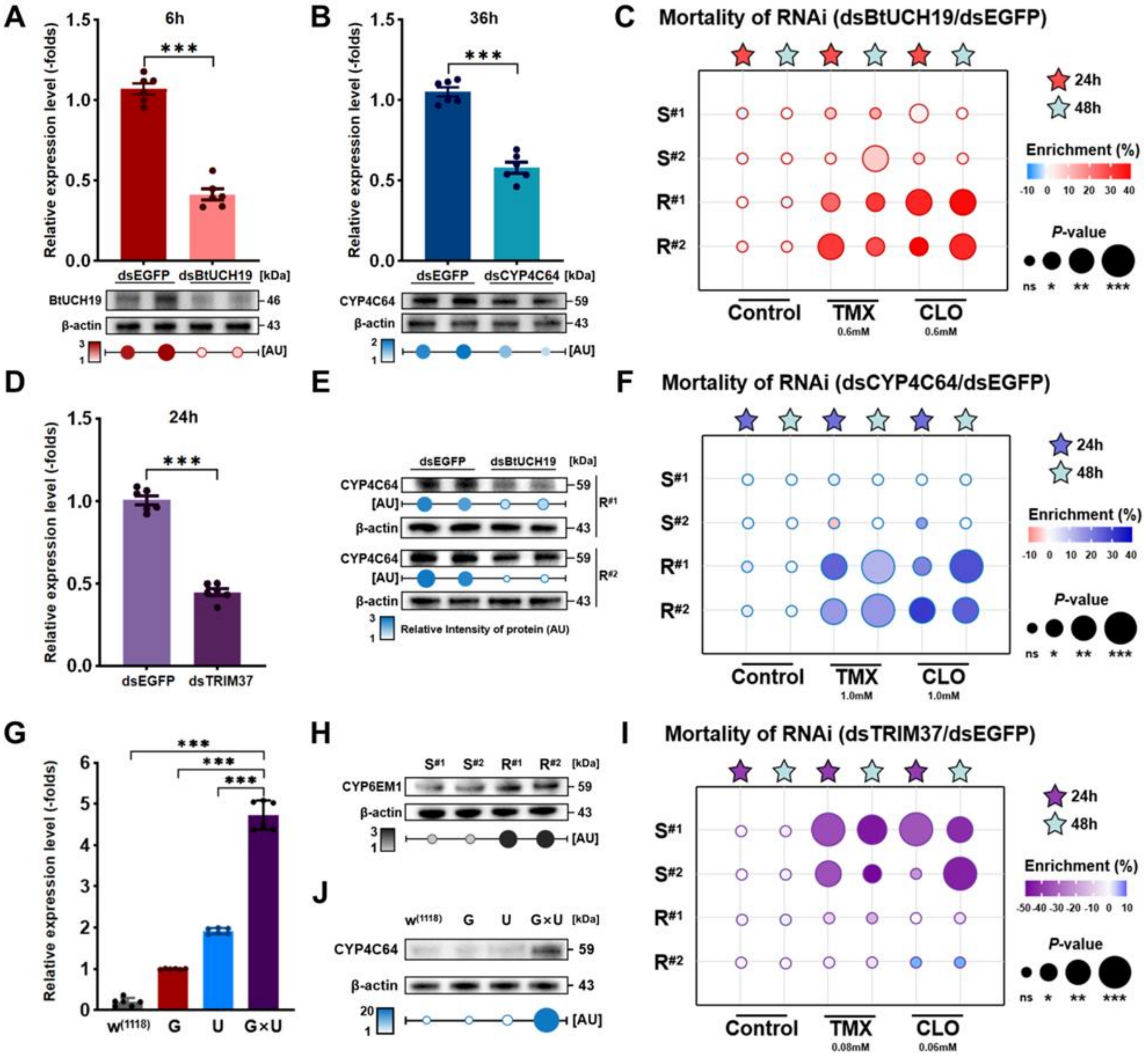
A
6h
Relative expression level (-folds)
dsEGFP dsBtUCH19 [kDa]
BtUCH19
β-actin
[AU]
B
36h
dsEGFP dsCYP4C64 [kDa]
CYP4C64
C
Mortality of RNAi (dsBtUCH19/dsEGFP)
24h
48h
Enrichment (%)
P-value
ns * ** ***
Control
TMX
0.6mM
CLO
0.6mM
D
24h
dsEGFP dsTRIM37
E
Relative Intensity of protein (AU)
F
Mortality of RNAi (dsCYP4C64/dsEGFP)
1.0mM
G
H
CYP6EM1
I
Mortality of RNAi (dsTRIM37/dsEGFP)
0.08mM
0.06mM
J

**Fig. S5. RNAi knockdown and transgenic *Drosophila* expressing *CYP4C64* provide functional evidence linking *BtUCH19*, *TRIM37*, and *CYP4C64* to neonicotinoid resistance in *B. tabaci*.** (A, B and D) Relative expression analyses of target genes and target proteins using qPCR and western blot after feeding on ds*BtUCH19* / 6h (A) and ds*CYP4C64* / 36h in resistant population $R^{\#2}$ (B), feeding on ds*TRIM37* / 24h with susceptible population $S^{\#2}$ (D), respectively. (C, F and I) Sensitivity of the experimental populations to thiamethoxam (TMX) and clothianidin (CLO) after RNAi knockdown of transgenes (*BtUCH19*, *CYP4C64* and *TRIM37*) under additional pesticide concentrations. (H) Detecting the protein level of CYP6EM1 among the four experimental populations. (E) Detecting the protein level of CYP4C64 in $R^{\#1}$ and $R^{\#2}$ after RNAi knockdown of *BtUCH19*. (G) Expression of *CYP4C64* in transgenic and parental *Drosophila* lines. (J) Detecting the protein levels of CYP4C64 in transgenic and parental *Drosophila* expressing CYP4C64, transgenic *Drosophila* expressing CYP4C64 compared to the wild-type line and the parental line that does not actively express the transgene. Note that all qPCR data are presented as the mean ± SEM of at least three independent experiments. All bioassay data (mean ± SEM) represent at least three biological replicates with approximately 60 adults in each replication and mortality was assessed 24 h and 48 h after first exposure to insecticide. For *B. tabaci*, newly emerged (1-day-old) adults were used in the RNAi experiments and whiteflies fed on ds*EGFP* were used as a control; For transgenic *Drosophila*, $w^{(1118)}$, wild type line; G, *Tub-GAL4/*$^{+}$ line; U, *UAS-CYP4C64/*$^{+}$ line; G×U, *UAS-CYP4C64/Tub-GAL4* line. All western blot analyses represent at least three biological replicates and *β*-actin was used as a loading control. AU, Relative Intensity of protein. Data were analyzed with ANOVA with Tukey's HSD post hoc test: ***$P < 0.001$.

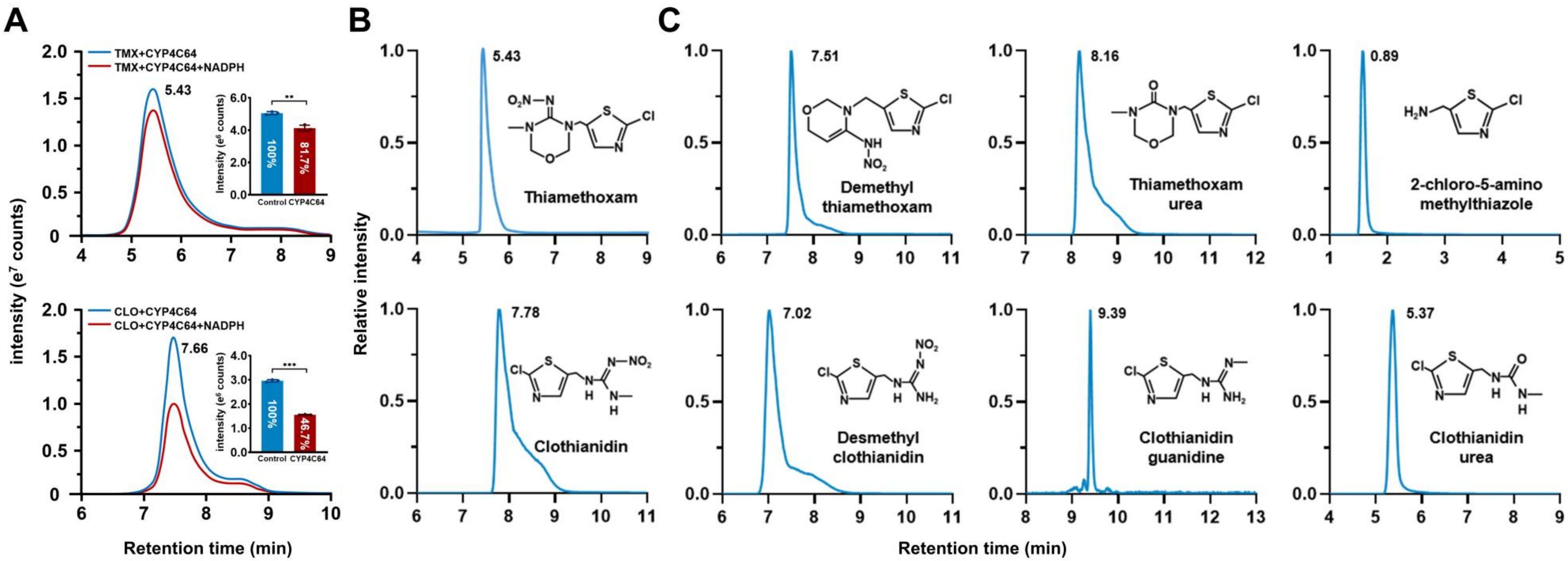


**Fig. S6. Analysis of Metabolites of Neonicotinoids Metabolised by CYP4C64.** (A) The residue of thiamethoxam (TMX, 50 pM) and clothianidin (CLO, 40 pM) are shown in the form of a column chart. (B and C) The retention time of neonicotinoid insecticides and their metabolites. (B) The retention times of TMX and CLO. (C) The retention times of demethyl-thiamethoxam, thiamethoxam-urea (TMX-Urea), 2-chloro-5-amino methylthiazole; Desmethyl-clothianidin, clothianidin-guanidine (CLO-Gdm), clothianidin-urea (CLO-Urea). Note that all UPLC-MS analyses data are presented from at least three independent experiments. Mass concentrations were calculated using relative standard curves, with microsomes incubated without NADPH serving as the control. All data were analyzed with ANOVA with Tukey's HSD post hoc test: **P* < 0.05, ***P* < 0.01 and ****P* < 0.001.

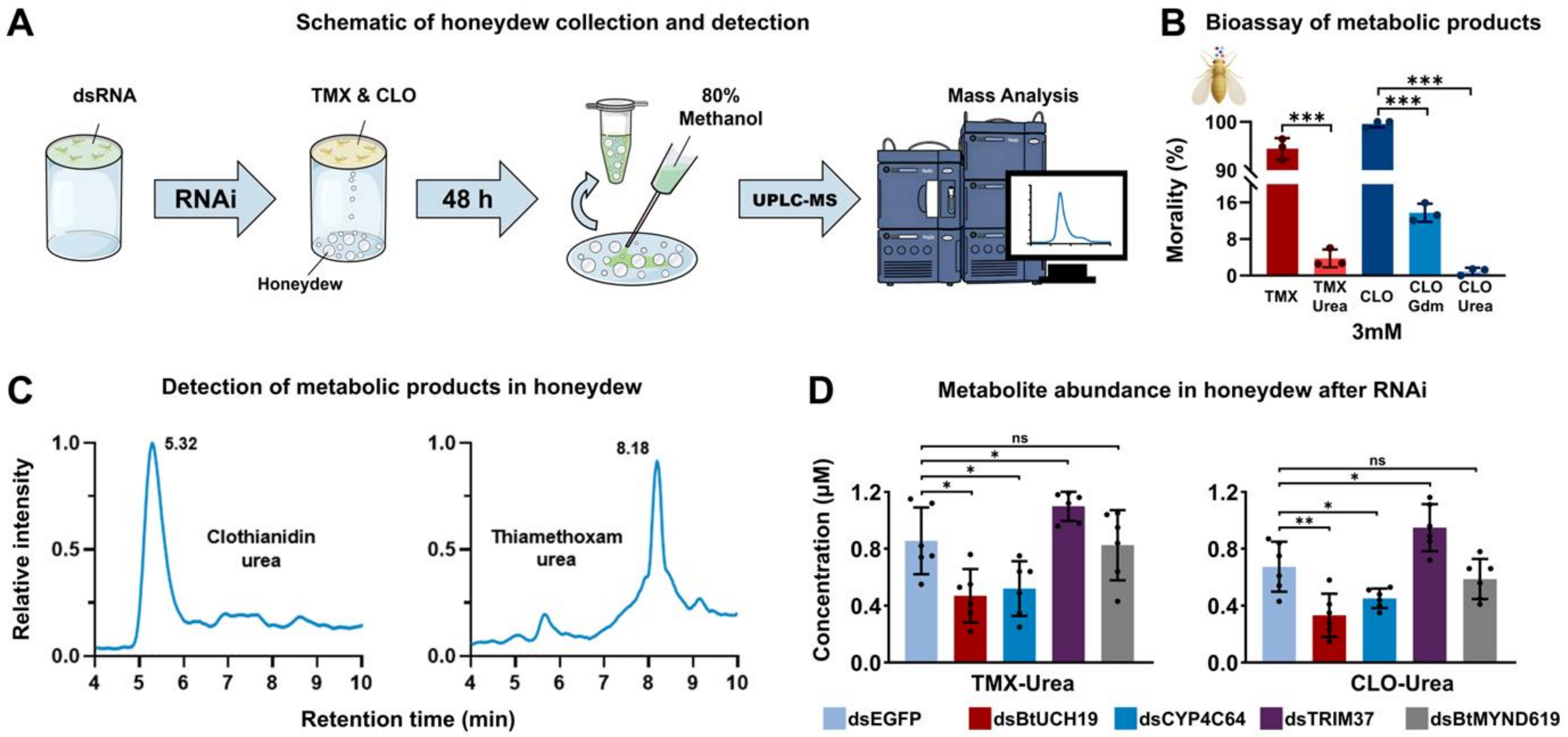


**Fig. S7. Toxicological Analysis of Metabolites of Neonicotinoids.** (A) Illustration of the whitefly honeydew collection setup and analysis. The honeydew collection device consists of a glass tube and a plastic bottom covered with tinfoil. The honeydew produced within 48 h was washed with 1 mL of 80% methanol aqueous solution and quantified. After filtration, the solution was quantitatively analyzed for metabolites using UPLC-MS. Every honeydew collection device includes 20 male and 60 female, with each test representing at least six replicates independently. (B) The sensitivity of $R^{\#2}$ to TMX, CLO, TMX-Urea, CLO-Gdm and CLO-Urea in 3 mM. (C) Detection of metabolites of neonicotinoids in population $R^{\#2}$ whitefly honeydew. (D) Feeding the diet solution containing TMX and CLO after interfering with *CYP4C64* / *BtUCH19* / *TRIM37* / *BtMYND619* in population $R^{\#2}$, then detecting the changes in the component contents (TMX-Urea and CLO-Urea) of honeydew. Note that all UPLC-MS analyses data are presented from at least three independent experiments. Mass concentrations were calculated using relative standard curves, with whiteflies fed on ds*EGFP* serving as the control. All bioassay data (mean ± SEM) represent at least three biological replicates with approximately 60 adults in each replication and mortality was assessed 48 h after first exposure to insecticide, newly emerged (1-day-old) adults were used in the RNAi experiments and whiteflies fed on ds*EGFP* serving as a control. All data were analyzed with ANOVA with Tukey's HSD post hoc test: *$P < 0.05$, **$P < 0.01$ and ***$P < 0.001$.

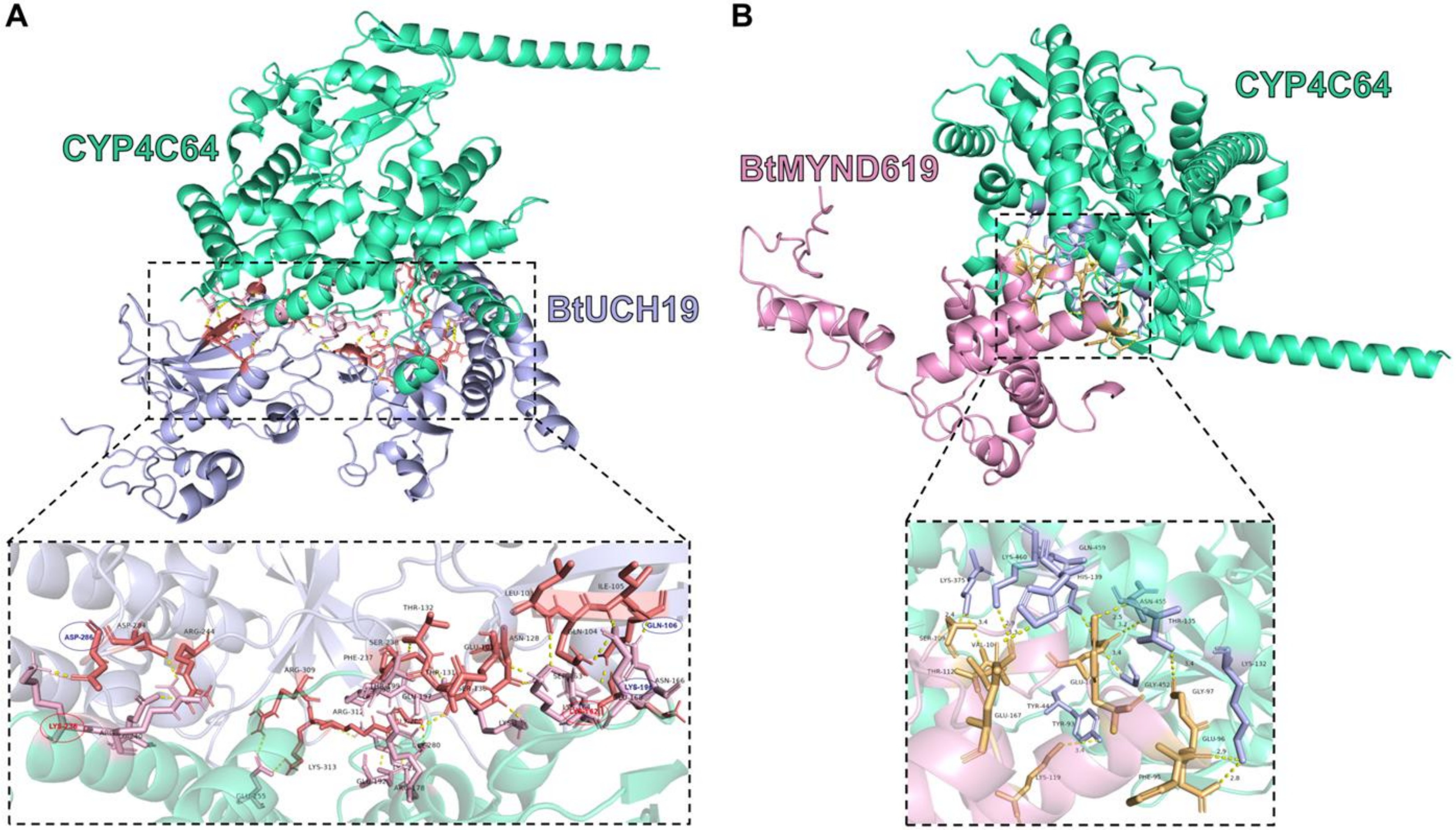


**Fig. S8. Prediction of CYP4C64 deubiquitination sites and protein docking analysis.** (A) Interconnection prediction of BtUCH19 (purple) and CYP4C64 (green). A hydrogen bond and salt bridge are formed between Asp-286 of BtUCH19 and Lys-236 of CYP4C64; Glu-106 and Lys-169 of BtUCH19 form hydrogen bonds with Lys-162 of CYP4C64. (B) Interconnection prediction of BtMYND619 (red) and CYP4C64 (green). Note that hydrogen bond (yellow lines) and salt bridge (red lines).

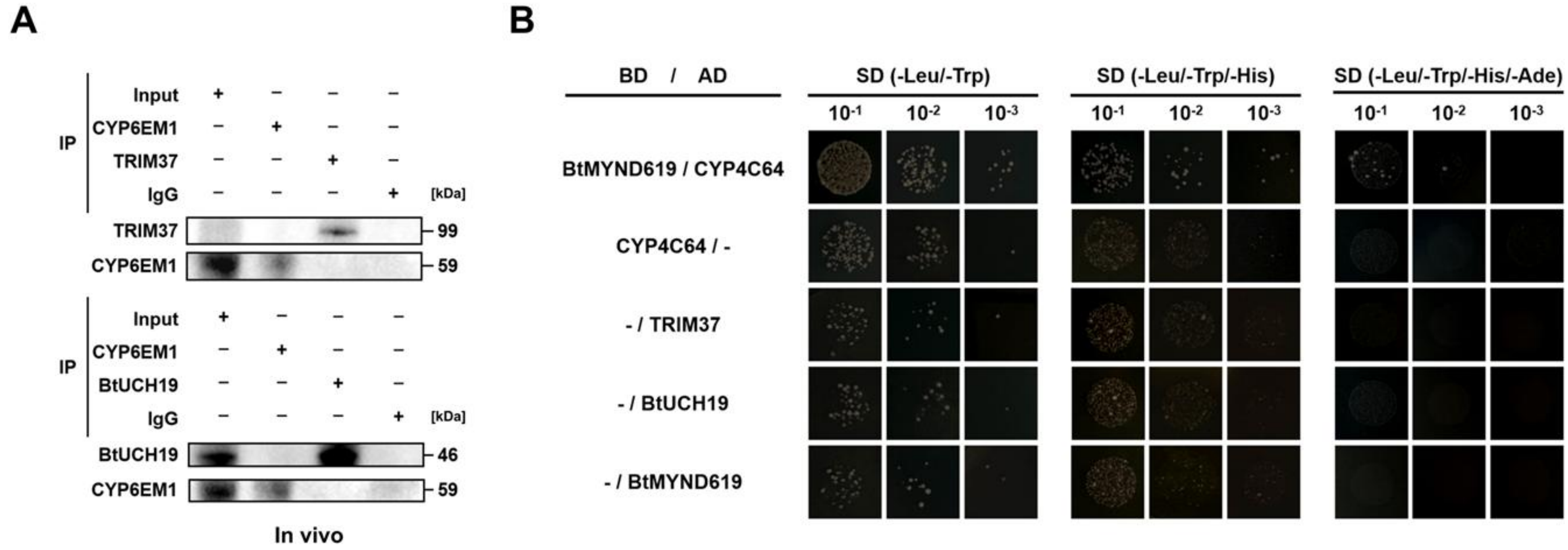


**Fig. S9. Verification of the specificity of BtUCH19 binding to CYP4C64.** Exclude false positives in experiments. (A) Immunoprecipitation (IP) analyses of interaction of BtUCH19 / TRIM37 with CYP6EM1 in the resistant population $R^{\#2}$. The extracted protein from whiteflies was incubated with the antibody of BtUCH19 / TRIM37 and CYP6EM1, followed by incubation with protein A / G-magnetic beads. Then the BtUCH19 / TRIM37 and CYP6EM1 proteins were detected by their corresponding antibodies in western blots. (B) Y2H assays showing the interaction between BtMYND619 and CYP4C64 *in vitro*. Exclude the possibility that BtUCH19 and TRIM37 do not undergo self-association, as well as the possibility that CYP4C64 does not undergo self-activation. Note that AD (Activation domain), BD (DNA binding domain), SD (Synthetic dextrose minimal medium), - (lacking), Ade (Adenine), His (Histidine), Leu (Leucine), Trp (Tryptophan). All western blot analyses represent at least three biological replicates, Input and IgG (immunoglobulin G) were used as loading controls.

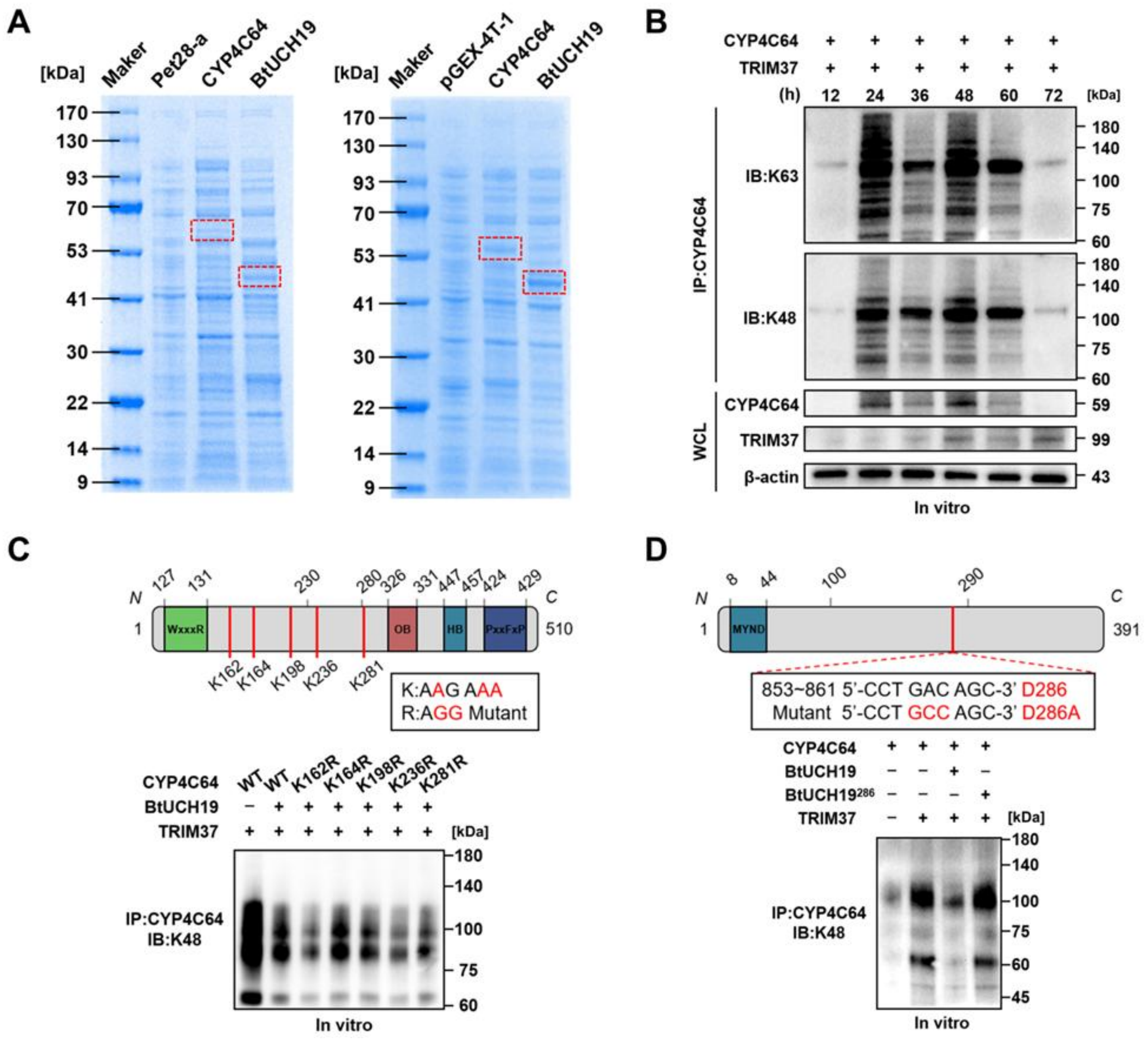

A
[kDa] Maker Pet28-a CYP4C64 BtUCH19
170 130 93 70 53 41 30 22 14 9
[kDa] Maker pGEX-4T-1 CYP4C64 BtUCH19
B
CYP4C64
TRIM37
(h) 12 24 36 48 60 72 [kDa]
IP:CYP4C64
IB:K63
IB:K48
WCL
CYP4C64 59
TRIM37 99
β-actin 43
In vitro
C
N 127 131 230 280 326 331 447 457 424 429 C
1 WxxxR OB HB PxxFxP 510
K162 K164 K198 K236 K281
K:AAG AAA
R:AGG Mutant
CYP4C64 WT WT K162R K164R K198R K236R K281R
BtUCH19
TRIM37 [kDa]
IP:CYP4C64
IB:K48
In vitro
D
N 8 44 100 290 C
1 MYND 391
853~861 5'-CCT GAC AGC-3' D286
Mutant 5'-CCT GCC AGC-3' D286A
CYP4C64
BtUCH19
BtUCH19286
TRIM37 [kDa]
IP:CYP4C64
IB:K48
In vitro

**Fig. S10. Characterisation of the binding site of BtUCH19 and CYP4C64 and their molecular interaction.** (A) Prokaryotic expression of the recombinant CYP4C64 and BtUCH19 protein in Escherichia coli as detected by SDS-PAGE precast gel. The recombinant Pet28a-CYP4C64 and Pet28a-BtUCH19 proteins were induced by 0.5 mM IPTG at 16 °C for 48 h. The recombinant pGEX-4T-1-CYP4C64 and pGEX-4T-1-BtUCH19 were induced by 0.5 mM IPTG at 23 °C for 24 h. The empty vector Pet28a and pGEX-4T-1 were used as controls. The red boxes indicate the in vitro-purified proteins after staining with InstantBlue Coomassie Protein Stain. (B) The effect of CYP4C64 ubiquitination level. The total protein was extracted from S2 cells after transfer for 12-72 h, then the protein was immunoprecipitated with anti-CYP4C64 and detected by anti-K63 / K48 in western blots. (C-D) IP and western blotting analyses of the ubiquitination (K48) of CYP4C64 in S2 cells for 24 h; Schematic diagrams of CYP4C64 and BtUCH19 mutant constructs are shown (above). The western bolts tested the reciprocal interaction after mutation of CYP4C64 and BtUCH19 in transfected S2 cells, as detected by the signal intensity of CYP4C64 obtained with K48 antibody (Bottom). BtUCH19, interaction with CYP4C64 wild-type or CYP4C64 mutants (C); CYP4C64 interacts with BtUCH19 wild-type or BtUCH19 mutants (D). Note that WxxxR: WxxxR motif; OB: Oxygen-binding motif; HB: Heme-binding motif; PxxFxP: PxxFxP motif; MYND: Zinc finger domain. All western blot analyses represent at least three biological replicates and $\beta$-actin and WCL (whole cell lysates) were used as loading controls. Cell lines in (C-E) were co-transfected to express TRIM37.

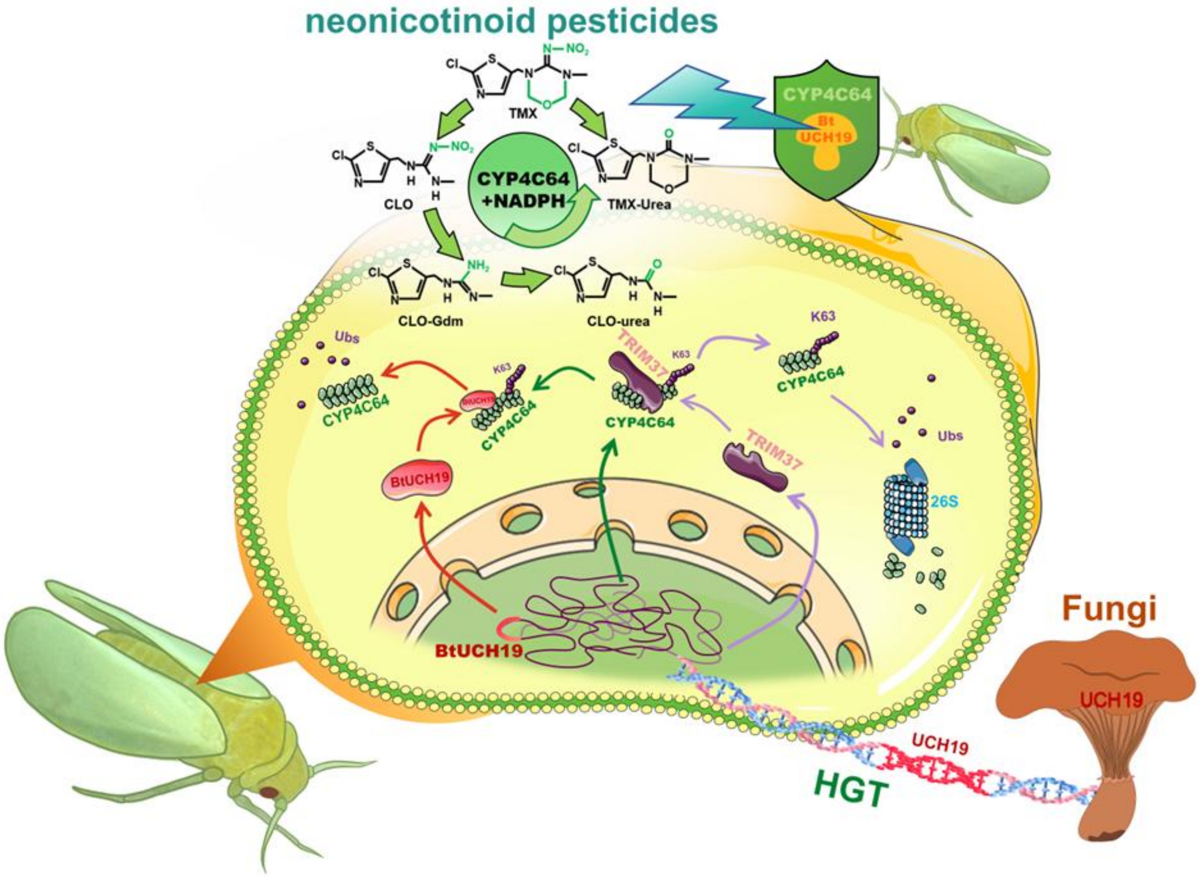


**Fig. S11. Model of CYP4C64 regulation by ubiquitination-deubiquitination in *B. tabaci.*** The deubiquitinating enzyme gene *BtUCH19* from fungi has been integrated into the genome of whiteflies through horizontal gene transfer and upregulated in neonicotinoid resistant strains of *B. tabaci*, then deubiquitinates the P450 CYP4C64. This ensures that CYP4C64 avoids ubiquitination by the E3 ubiquitin ligase TRIM37 and subsequent degradation via the 26S proteasome system. Specifically, BtUCH19 prevents k63-linked ubiquitination of *CYP4C64* at Lys162 and Lys236, maintaining the stability of CYP4C64 protein and preventing it from being degraded. This promotes the metabolism of thiamethoxam and clothianidin into thiamethoxam-urea and clothianidin-urea by CYP4C64.

**Table. S1.** Origin of the experimental strains of *B. tabaci* used in this study

| Population | Sampling location | Host plant | Collection date | Whitefly composition | Population type |
|---|---|---|---|---|---|
| $S^{\#1}$ | Hangzhou, Zhejiang (30˚20′ N, 12˚38′ E) | Tomato | 2011.08 | MED / Q | Laboratory sample |
| $S^{\#2}$ | Nanning, Guangxi (22˚80′ N, 108˚20′ E) | Eggplant | 2021.07 | MED / Q | Field-collected sample |
| $R^{\#1}$ | Yueyang, Hunan (29˚21′ N, 113˚7′ E) | Pepper | 2023.03 | MED / Q | Field-collected sample |
| $R^{\#2}$ | Sanya, Hainan (18˚15′ N, 109˚30′ E) | Eggplant | 2023.04 | MED / Q | Field-collected sample |

**Table. S2.** Resistance ratios ($RR_{50}$) of the experimental populations of *B. tabaci* to two neonicotinoid insecticides

| Insecticides | Populations | N [a] | Slop (±SE) | $LC_{50}$ ($mg \cdot L^{-1}$) (95% FL [b]) | $x^2$ [c] | $RR_{50}$ [d] |
|---|---|---|---|---|---|---|
| Thiamethoxam (TMX) | $S^{\#1}$ | 389 | 2.75 (±0.12) | 1.02 (0.65-2.03) | 2.11 | 1.92 |
| | $S^{\#2}$ | 456 | 1.50 (±0.16) | 0.53 (0.40-0.67) | 1.28 | 1.00 |
| | $R^{\#1}$ | 443 | 1.46 (±0.19) | 169.27(100.69-452.77) | 4.50 | 319.38 |
| | $R^{\#2}$ | 434 | 1.50 (±0.30) | 127.58 (88.94-259.00) | 0.98 | 240.72 |
| Clothianidin (CLO) | $S^{\#1}$ | 359 | 2.22 (±0.15) | 1.44 (0.82-3.88) | 2.29 | 1.58 |
| | $S^{\#2}$ | 554 | 1.23 (±0.19) | 0.72 (0.42-1.10) | 4.01 | 1.00 |
| | $R^{\#1}$ | 413 | 2.20 (±0.28) | 139.96 (102.03-255.61) | 8.14 | 194.39 |
| | $R^{\#2}$ | 435 | 3.57 (±0.37) | 103.94 (82.76-137.38) | 7.11 | 144.36 |

[a] N, Number of adult whiteflies used in bioassay.
[b] FL, Fiducial limit.
[c] $x^2$, Chi-square.
[d] $RR_{50}$, Resistance ratio = $LC_{50}$ (test population) / $LC_{50}$ (the most susceptible population).
SE, Standard error; R, Resistant population; S, Susceptible population.

**Table. S3.** Information on key genes included in this study

| Gene Name [a] | Gene summary | Functions | Ref. |
|---|---|---|---|
| CYP4C64 (GenBank JX144366) | Cytochrome P450 detoxifying enzyme | Detoxifying enzymes | (13) |
| TRIM37 (GenBank XM_019044747) | E3 ubiquitin ligase of *Bemisia. tabaci* | Protein degradation | Hu |
| BtUCH19 (GenBank XM_072302559) | Ubiquitin carboxyl-terminal hydrolase of *Bemisia. tabaci* | Protein stability | - |
| BtCPR (GenBank XM_019058755.2) | NADPH-Cytochrome P450 reductase of *Bemisia. tabaci* | Electron donor | (14) |
| BtMYND619 (GenBank XR_011900330.1) | Ubiquitin carboxyl-terminal hydrolase of *Bemisia. tabaci* | Protein stability | - |
| EF-1α (GenBank EE600682) | Housekeeping genes of *Bemisia. tabaci* | Eukaryotic translation elongation factor | (2) |
| RPL29 (GenBank EE596314) | Housekeeping genes of *Bemisia. tabaci* | Ribosomal protein | (2) |
| RPL11 (GenBank NM_001299685) | Housekeeping genes of *Drosophila. melanogaster* | Ribosomal protein | (15) |

[a] Note that more detailed information can be found on the NCBI: pubmed.ncbi.nlm.nih.gov

**Table. S4.** Primers used in this study

| Usage [a] | Gene | Primer name [b] | Primer sequence (5′-3′) | Product size (bp) | Tm (°C) | Amplification efficiency (%) |
|---|---|---|---|---|---|---|
| qPCR | CYP4C64 | qC64-F | TCGGATTACGTCAGAGCTATTTAC | 138 | 62 | 100 |
| | | qC64-R | GTGGAGCACGCTTAGACA | | 63 | |
| | TRIM37 | q37-F | TGAGGAGTTTGCTGGGTTCT | 147 | 61 | 99 |
| | | q37-R | TTTGTCCGGATTCCTTTTTG | | 61 | |
| | BtUCH19 | q19-F | CGGAAGTACCCTCCAAACCC | 54 | 61 | 98 |
| | | q19-R | TGTCAGGATCGGCGCATAAA | | 61 | |
| | BtMYND619 | q619-F | TGTTGAATATCGGTGCGCTACT | 100 | 59 | 99 |
| | | q619-R | ACACTCACGAGAGCAGTAG | | 58 | |
| | EF-1α | EF-F | TAGCCTTGTGCCAATTTCCG | 110 | 60 | 103 |
| | | EF-R | CCTTCAGCATTACCGTCC | | 60 | |
| | RPL29 | RP-F | TCGGAAAATTACCGTGAG | 144 | 60 | 101 |
| | | RP-R | GAACTTGTGATCTACTCCTCTCGTG | | 60 | |
| | RPL11 | RP11-F | GTATTCGCCGTAACGAGAAGAT | 146 | 60 | 99 |
| | | RP11-R | GATGCCGAAACCGAAGTTGC | | 62 | |

[a] qPCR: Quantitative reverse-transcription polymerase chain reaction.
[b] F, Forward primer; R, Reverse primer.

| Usage | Gene | Primer name | Primer sequence (5′-3′) | Product size (bp) | Tm (°C) | Database |
|---|---|---|---|---|---|---|
| Gene cloning | CYP4C64 | C64-F | TTTCTCGGGGTCAGCCTTGC | 1533 | 62 | Bta12387 |
| | | C64-R | CTAAAATAGTAGGTAAAGTAATCGC | | 63 | |
| | TRIM37 | Ub3-F | TTTGATAGCGGGCTCTCGTC | 2784 | 60 | Bta11294 |
| | | Ub3-R | AGTGCTGCGTCATTGAAGAGG | | 60 | |
| | BtUCH19 | U19-F | CAGTGAACTACAGTCTGCGTTTC | 1659 | 62 | Bta10703 |
| | | U19-R | CCTTGCACCTAAGAGGCTAC | | 61 | |
| | BtMYND619 | 619-F | ATGTCCTCGAATGTTGAATATCGGTG | 600 | 59 | Bta11249 |
| | | 619-R | TCACTCGCTGTCTGAGTCAGG | | 60 | |
| | BtCPR | CPR-F | ATGAAATACCTGCTGCCGACCG | 2048 | 61 | Bta14494 |
| | | CPR-R | GCTTGACTGTCTGTCATGTTACCT | | 59 | |

Note that more detailed information can be found on the Whitefly genome database: whiteflygenomics.org

| Usage | Gene | Primer name | Primer sequence (5′-3′) | Product size (bp) |
|---|---|---|---|---|
| RNAi | CYP4C64 | dsC64-F | TAATACGACTCACTATAGGGAGAATTTGTGAGACCGCTATGGG | 421 |
| | | dsC64-R | TAATACGACTCACTATAGGGAGATGGTTGTATCGTGTCCCTCA | |
| | TRIM37 | dsUb3-F | TAATACGACTCACTATAGGAATGCTGTAAAAGGTAGCTCTTCA | 374 |
| | | dsUb3-R | TAATACGACTCACTATAGGACTATCACTAAATGTTGGACTGGC | |
| | BtUCH19 | dsU19-F | TAATACGACTCACTATAGGGAGACCTGTGCTCAGACTATGCCA | 500 |
| | | dsU19-R | TAATACGACTCACTATAGGGAGAGATCCCGGTCACCAACACAA | |
| | BtMYND619 | ds619-F | TAATACGACTCACTATAGGGAGACTTTGCAGTTCTTGATGGCA | 310 |
| | | ds619-R | TAATACGACTCACTATAGGGAGAACCGATAAAACTCCAGCCCT | |
| | EGFP | dsEGFP-F | TAATACGACTCACTATAGGGAGACAGTGCTTCAGCCGCTAC | 288 |
| | | dsEGFP-R | TAATACGACTCACTATAGGGAGAGTTCACCTTGATGCCGTTC | |
| Vector linearization | pAc5.1b | pAc-F | GACACAAAGCCGCTCCATCAG | 252 |
| | | pAc-R | CCTTAGAAGGCACAGTCGAGGC | |
| | pGL4.26 | pGL-F | TAGCAAAATAGGCTGTCCCCAG | 282 |
| | | pGL-R | CGTCGGTAAAGGCGATGGTG | |

**Table. S5.** Cell lines used in this study

| Cell line | Source | Cat# |
|---|---|---|
| Schneider 2: S2 | Thermo Fisher | R69007 |
| Spodoptera frugiperda: Sf9 | Invitrogen | B82501 |
| High Five: H5 | Invitrogen | B85502 |

**Table. S6.** List of primer sequences used in genome fragment cloning

| Location | Gene | Primer name | Primer sequence (5′-3′) | Tm (°C) | Product size (bp) |
|---|---|---|---|---|---|
| Scaffold467 Genome fragment cloning | 1775192-1779400 | S-F1 | CATTGGTGGTGGTGGAGCTG | 61 | 4189 |
| | | S-R1 | AGTCTGAGCACAGGAGTCGG | 61 | |
| | 1780143-1782742 | S-F2 | CCAACGTTGGCAAGCAGGAT | 61 | 2424 |
| | | S-R2 | GTCAGGCAGTGTTGTCGCAT | 60 | |
| | 1782651-1785126 | S-F3 | GCGGGAAGCATAGCCAATCG | 60 | 2301 |
| | | S-R3 | CGCCTACAAACCTGACAGGG | 59 | |
| | 1784994-1787815 | S-F4 | TCAAGCGCCGTGGTATGCTG | 62 | 2580 |
| | | S-R4 | TTCACGGCTGTAATGAGTGCC | 61 | |
| | 1787643-1790240 | S-F5 | CGTAGTCTGGAGCCGGCTTT | 61 | 2693 |
| | | S-R5 | CGTGGTTTTTCTTCTCTGTGCG | 60 | |
| | 1789786-1792592 | S-F6 | GTGCTGCCGGCTATAGAAGC | 61 | 2683 |
| | | S-R6 | GTTGGCGACGTTGCAAACTT | 60 | |

**Table. S7.** List of mutant sequences used in this study

| Vector | Genes | Name | Original sequences (5′-3′) | Mutant sequences (5′-3′) | Positions |
|---|---|---|---|---|---|
| pAC5.1b | CYP4C64 | C64 (WT) | — | — | 2615 / 4147 |
| | | $C64^{162}$ | AAG | AGG | 3098 / 3100 |
| | | $C64^{164}$ | AAA | AGG | 3104 / 3106 |
| | | $C64^{198}$ | AAA | AGG | 3206 / 3208 |
| | | $C64^{236}$ | AAA | AGG | 3320 / 3322 |
| | | $C64^{281}$ | AAA | AGG | 3455 / 3457 |
| | BtUCH19 | U19 (WT) | — | — | 2615 / 3790 |
| | | $U19^{286}$ | GAC | GCC | 3470 / 3472 |
| | TRIM37 | Ub3 | — | — | 2615 / 5239 |
| PGADT7(AD) | CYP4C64 | AD-C64 | — | — | 1972 / 3504 |
| | | $AD\text{-}C64^{162}$ | AAG | AGG | 2455 / 2457 |
| | | $AD\text{-}C64^{164}$ | AAA | AGG | 2461 / 2463 |
| | | $AD\text{-}C64^{198}$ | AAA | AGG | 2563 / 2565 |
| | | $AD\text{-}C64^{236}$ | AAA | AGG | 2677 / 2679 |
| | | $AD\text{-}C64^{281}$ | AAA | AGG | 2812 / 2814 |
| | BtUCH19 | AD-U19 | — | — | 1972 / 3630 |
| | TRIM37 | AD-Ub3 | — | — | 1972 / 4755 |
| | BtMYND619 | AD-619 | — | — | 1972 / 1571 |
| PGBKT7(BD) | CYP4C64 | BD-C64 | — | — | 1284 / 2816 |
| | BtUCH19 | BD-U19 | — | — | 1284 / 2459 |
| | TRIM37 | BD-Ub3 | — | — | 1284 / 3908 |
| | BtMYND619 | BD-619 | — | — | 1284 / 1883 |

| Vector | Genes | Name | Product size (bp) | Kilodalton (kDa) | Positions |
|---|---|---|---|---|---|
| pGEX-4T | CYP4C64 | GST-C64 1-510 | 1533 | 59.1 | 936 / 2468 |
| | | GST-C64 230-510 | 846 | 32.4 | 936 / 1781 |
| | | GST-C64 1-280 | 843 | 33.4 | 936 / 1778 |
| | | GST-C64 1-229 | 690 | 26.7 | 936 / 1625 |
| | | GST-C64 281-510 | 693 | 26.7 | 936 / 1628 |
| | BtUCH19 | GST-U19 1-391 | 1176 | 44.9 | 936 / 2111 |
| | | GST-U19 1-290 | 873 | 33.1 | 936 / 1808 |
| | | GST-U19 100-391 | 897 | 33.5 | 936 / 1814 |
| | | GST-U19 291-391 | 306 | 11.9 | 936 / 1241 |
| | | GST-U19 1-99 | 303 | 11.6 | 936 / 1238 |
| | GST | GST | 657 | 25.45 | — |
| pET-28a | CYP4C64 | His-C64 1-510 | 1533 | 59.1 | 5131 / 6663 |
| | BtUCH19 | His-U19 1-391 | 1176 | 44.9 | 5131 / 6306 |
| | His | His | — | — | — |
| pFast Bac 1 | CYP4C64 | pFast Bac-C64 | 1533 | 55.9 | 4060 / 5592 |
| | BtCPR | pFast Bac-CPR | 2048 | 76.7 | 4060 / 6107 |
| pJFRC28 | CYP4C64 | p10-C64 | 1533 | 55.9 | 7093 / 8625 |
| | BtCPR | p10-CPR | 2048 | 76.7 | 7093 / 9140 |

566

**Table. S8.** Details of antibodies used in this study

| Antibody | Availability | Peptides site | Source | Ref. |
|---|---|---|---|---|
| BtUCH19 | synthesis compound | KRQFNQVADEPNPFD （Lys376-Asp390） | rabbit | — |
| CYP4C64 | synthesis compound | NKRIQLVRTMN (Asn25-Asn35) | rabbit | (13) |
| TRIM37 | commercial product | (#PA5-30094, Invitrogen) | rabbit | Hu |
| CYP402C9 | synthesis compound | QKYEVRPHSHTKQMD (Gln479-Asp493） | rabbit | (4) |
| CYP6EM1 | synthesis compound | QNNFTYWRKRGVPYV (Gln24-Val38) | rabbit | (16) |
| CYP6CX3 | synthesis compound | FHDRGPEPDEKVDKLSGN (Phe44-Asp61) | rabbit | (17) |
| CYP306A1 | synthesis compound | SPNTPEELQNALDGC (Ser470-Cys483) | rabbit | (18) |
| CYP6DW3 | synthesis compound | KKRYEYWQKLA (Lys25-Ala35) | rabbit | (19) |
| CYP4G68 | synthesis compound | GNAYEFMGDSNKIFENIC (Gly68-Cys84) | rabbit | (20) |
| CYP4G129 | synthesis compound | IKKKKEDYRLRKNGLID (Ile266-Asp282) | rabbit | (4) |
| CYP6CM1 | synthesis compound | TTPKTPKKITFDTN (Tyr486-Asn500) | rabbit | (21) |
| GSTS1 | synthesis compound | GTMSYFAKIDLV ( Gly161-Val184) | rabbit | (22) |
| GSTS2 | synthesis compound | VEFIDDRVNFVED (Val27-Asp39) | rabbit | (22) |
| GSTS4 | synthesis compound | ANYYYDPDEESKRRKY (Ala103-Tyr118) | rabbit | (22) |
| GSTZ1 | synthesis compound | WKKDEHLTPEYRAI (Trp40-Ile53) | rabbit | (22) |
| UGT354A1 | synthesis compound | RIFRSIFSTETKQKT (Arg498-Thr512) | rabbit | (23) |
| UGT356B1 | synthesis compound | KSFKNFEHIAISADTSC (Lys64-Ser79) | rabbit | (23) |
| *β*-actin | commercial product | (#ab8227, Abcam) | rabbit | — |
| His-Tag | commercial product | (#ab18184, Abcam) | rabbit | — |
| GST-Tag | commercial product | (#ab9085, Abcam) | rabbit | — |
| K48 | commercial product | (#PTM-7228, PTM BIO) | rabbit | — |
| K63 | commercial product | (#PTM-7205, PTM BIO) | rabbit | — |
| IgG-HRP | commercial product | (#CW0103S, CWBIO) | goat anti-rabbit | — |

**Table. S9.** Kinetics of CYP4C64-catalysed neonicotinoids

| **Substrate** | **$V_{max}$ [a] (95% FL [b])** | **$K_m$ [c] (95% FL)** | **$K_{cat}$ [a]** | **N [d]** | **Df [e]** | **$R^2$ [f]** |
|---|---|---|---|---|---|---|
| Thiamethoxam | 2.42 (0.23-0.64) | 17.16 (12.80-22.92) | 0.14 | 24 | 22 | 0.951 |
| Clothianidin | 4.40 (4.01-4.86) | 11.69 (8.54-16.00) | 0.38 | 24 | 22 | 0.944 |

[a] pmol / min / pmol $^{P450}$.
[b] FL, Fiducial limit.
[c] μmol.
[d] *N*, Number of testing object.
[e] Df, Degrees of freedom.
[f] $R^2$, Coefficient of determination.

$V_{max}$: Maximum metabolic rate; $K_m$: Michaelis constant; $K_{cat}$: Catalytic constant ($V_{max}$ / $K_m$).

**Table. S10.** Interface parameters between CYP4C64 and BtUCH19 / BtMYND619

| Protein | Interface area (Å) | $\Delta^iG$ (Kcal/mol) | $\Delta^iG$ (*P*-value) | $N_{HB}$ | $N_{SB}$ | $N_{DS}$ | CSS |
|---|---|---|---|---|---|---|---|
| BtUCH19 | 1933.4 | -10.9 | 0.557 | 15 | 12 | 0 | 0.000 |
| BtMYND619 | 1003.7 | -1.0 | 0.729 | 9 | 3 | 0 | 0.000 |

Interface area: Area of protein interaction interface.
$\Delta^iG$: Solvation free energy gain.
N: Number of chemical bonds.
$_{HB}$: Hydrogen bonds.
$_{SB}$: Salt bridges.
$_{DS}$: Disulfide bonds.
CSS: Complexation significance score.

Note that all parameters were simulated using PDBePISA (https://www.ebi.ac.uk/msd-srv/prot_int/).

**Table. S11.** Experimental standard products and UPLC-MS parameters

| Chemical formula CAS [a] | Skeletal formula | CYP4C64 -NADPH | CYP4C64 +NADPH | Source formulation [b] | Retention time (min) | Transitions (m/z) | CV (V) [c] | CE (V) [d] | DT (ms) [e] |
|---|---|---|---|---|---|---|---|---|---|
| Thiamethoxam<br>$C_8H_{10}ClN_5O_3S$<br>153719-23-4 | 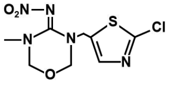 | + | - | Shanghai Yuanye Bio-Tech Co., Ltd.<br>99% TC | 5.43 | 292.0-181.0 | 22 | 20 | 0.065 |
| | | | | | | 292.0-211.0 | 22 | 10 | 0.065 |
| 2-chloro-5-amino-methylthiazole<br>$C_4H_5ClN_2S$<br>120740-08-1 | 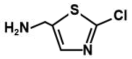 | - | - | Dr. Ehrenstorfer GmbH.<br>98% TC | 0.89 | 149.0-59.0 | 18 | 24 | 0.065 |
| | | | | | | 149.0-119.9 | 18 | 12 | 0.065 |
| Thiamethoxam-urea<br>$C_8H_{10}ClN_3O_2S$<br>902493-06-5 | 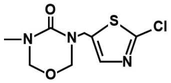 | - | + | Dr. Ehrenstorfer GmbH.<br>99% TC | 8.16 | 248.0-43.9 | 12 | 10 | 0.065 |
| | | | | | | 248.0-174.9 | 12 | 16 | 0.065 |
| Demethyl-thiamethoxam<br>$C_7H_8ClN_5O_3S$<br>171103-04-1 | 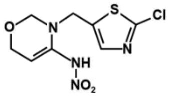 | - | - | Alta Scientific Co., Ltd.<br>98% TC | 7.51 | 278.0-131.9 | 30 | 10 | 0.065 |
| | | | | | | 278.0-196.9 | 30 | 10 | 0.065 |
| Clothianidin<br>$C_6H_8ClN_5O_2S$<br>210880-92-5 | 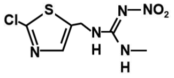 | + | + | Shanghai Yuanye Bio-Tech Co., Ltd.<br>98% TC | 7.78 | 245.0-132.0 | 30 | 12 | 0.078 |
| | | | | | | 245.0-168.9 | 30 | 10 | 0.078 |
| Desmethyl-clothianidin<br>$C_5H_6ClN_5O_2S$<br>135018-15-4 | 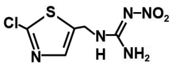 | - | - | Tan-Mo Technology Co., Ltd.<br>98% TC | 7.02 | 236.0-112.9 | 22 | 26 | 0.078 |
| | | | | | | 236.0-131.9 | 22 | 12 | 0.078 |
| Clothianidin-urea<br>$C_6H_8ClN_3OS$<br>634192-72-6 | 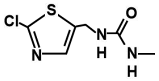 | - | + | Dr. Ehrenstorfer GmbH<br>99% TC | 5.37 | 206.0-112.9 | 4 | 16 | 0.078 |
| | | | | | | 206.0-132.0 | 4 | 12 | 0.078 |
| Clothianidin-guanidine hydrochloride<br>$C_6H_9ClN_4S \cdot HCl$<br>939773-18-9 | 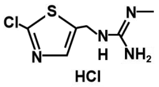 | - | + | Dr. Ehrenstorfer GmbH<br>86% TC | 9.39 | 204.99-119.94 | 80 | 12 | 0.078 |
| | | | | | | 204.99-177.01 | 80 | 4 | 0.078 |

[a] Source of the compounds and CAS: pubchem.ncbi.nlm.nih.gov
[b] TC: Technical compounds.
[c] CV: Cone voltage.
[d] CE: Collision energy.
[e] DT: Dwell time.
